# Global proteome and phosphoproteome alterations in third generation EGFR TKI resistance reveal drug targets to circumvent resistance

**DOI:** 10.1101/2020.07.04.187617

**Authors:** Xu Zhang, Tapan K. Maity, Karen E. Ross, Yue Qi, Constance M. Cultraro, Shaojian Gao, Khoa Dang Nguyen, Julie Cowart, Fatos Kirkali, Cathy Wu, Udayan Guha

**Affiliations:** Thoracic and GI Malignancies Branch, CCR, NCI, NIH, Bethesda, MD; Dept. of Biochemistry and Molecular & Cellular Biology, Georgetown University Medical Center, Washington DC; Center for Bioinformatics and Computational Biology, University of Delaware, Newark, DE

## Abstract

Lung cancer is the leading cause of cancer mortality worldwide. The treatment of lung cancer patients harboring mutant EGFR with orally administered EGFR TKIs has been a paradigm shift. Osimertinib and rociletinib are 3^rd^ generation irreversible EGFR TKIs targeting the EGFR T790M mutation. Osimertinib is the current standard care for patients with EGFR mutations due to increased efficacy, lower side effects, and enhanced brain penetrance. Unfortunately, all patients develop resistance. Genomic approaches have primarily been used to interrogate resistance mechanisms. Here, we have characterized the proteome and phosphoproteome of a series of isogenic EGFR mutant lung adenocarcinoma cell lines that are either sensitive or resistant to these drugs. To our knowledge, this is the most comprehensive proteomic dataset resource to date to investigate 3^rd^ generation EGFR TKI resistance in lung adenocarcinoma. We have interrogated this unbiased global quantitative mass spectrometry dataset to uncover alterations in signaling pathways, reveal a proteomic signature of epithelial mesenchymal transition (EMT) and identify kinases and phosphatases with altered expression and phosphorylation in TKI resistant cells. Decreased tyrosine phosphorylation of key sites in the phosphatase SHP2 suggests its inhibition resulting in inhibition of RAS/MAPK and activation of PI3K/AKT pathways. Furthermore, we performed anticorrelation analyses of this phosphoproteomic dataset with the published drug-induced P100 phosphoproteomic datasets from the Library of Integrated Network-Based Cellular Signatures (LINCS) program to predict drugs with the potential to overcome EGFR TKI resistance. We identified dactolisib, a PI3K/mTOR inhibitor, which in combination with osimertinib overcomes resistance both *in vitro* and *in vivo*.

**One Sentence Summary:** Global quantitative proteome and phosphoproteome analyses to examine altered signaling pathways in isogenic 3^rd^ generation EGFR TKI sensitive and resistant cells.

## Introduction

Lung cancer continues to be the leading cause of cancer mortality in the world (*1*). Many lung adenocarcinoma patients with activating epidermal growth factor receptor (EGFR) mutations initially respond dramaticlly to the first- or second-generation EGFR tyrosine kinase inhibitors (TKIs). However, they eventually develop resistance. The most common mechanism of acquired resistance is the EGFR T790M gatekeeper site residue mutation (*2*). Osimertinib, a third generation irreversible EGFR TKI has been approved by the FDA to treat patients harboring the EGFR T790M mutation who have developed resistance to first- and second-generation EGFR TKIs (*3*). Recently, osimertinib was also approved for the front-line treatment of patients harboring EGFR mutations (*4*). Rociletinib is another irreversible inhibitor targeting the EGFR T790M mutation, which has minimal activity against wild-type EGFR. Both drugs have therapeutic benefits and have demonstrated activity in tumors with T790M-mediated resistance to other EGFR tyrosine kinase inhibitors (*5, 6*). Further development of rociletinib was ceased in 2016 due to less than expected efficacy, poor brain penetration leading to tumor progression in brain tissues and off-target effects on IGFR activation leading to hyperglycemia (*7, 8*).

Although 3^rd^-generation TKIs provide clinical benefit to most patients with EGFR mutations, some patients, demonstrating primary resistance, still do not respond to these inhibitors. Complete responses are rare, and all patients eventually develop resistance, suggesting primary and acquired resistance mechanisms decrease the efficacy of the drugs (*9, 10*). Genomic approaches have been used primarily to interrogate osimertinib resistance mechanisms (*9, 11-15*). Several mechanisms of osimertinib resistance have been identified (*16*), including novel second site EGFR mutations, activated bypass pathways such as MET amplification, HER2 amplification, RAS mutations, BRAF mutations, PIK3CA mutations, and novel fusion events (*17*). However, the resistance mechanism is complex and still not fully understood.

Previously, we have used SILAC-based quantitative phosphoproteomics to identify the global dynamic modification which occur upon treatment of TKI-sensitive and -resistant lung adenocarcinoma cells with the 1^st^ and 2^nd^ generation EGFR TKIs, erlotinib and afatinib. Utilizing this strategy, we identified the targets of mutant EGFR signaling pathways responsible for TKI resistance, and possible off-target effects of the drugs (*18, 19*). In this study, we employed SILAC-based quantitative mass spectrometry to characterize alterations in the proteome and phosphoproteome which occur upon acquired resistance and sought to identify novel mechanisms of resistance to the third generation EGFR TKIs, osimertinib and rociletinib. To our knowledge, this is the most comprehensive 3^rd^ generation EGFR TKI resistant proteome and phosphoproteome analysis resource available to date.

## Results

### Quantitative mass spectrometry to identify and quantify the global proteome and phosphoproteome

H1975, an EGFR-L858R/T790M mutant 3^rd^ generation EGFR TKI-sensitive cell line, and the corresponding isogenic osimertinib (AZR3 and AZR4) or rociletinib (COR1 and COR10) resistant cell lines were used for 3-state SILAC experiments to characterize the proteome and phosphoproteome by quantitative mass spectrometry. Cells were cultured in medium containing light, medium, and heavy labeled amino acids and maintained in the drug until 3 days before the experiment. Cells were subsequently treated with either DMSO or the 3^rd^ generation EGFR TKIs, osimertinib or rociletinib (**Fig. 1A**). Both proteome and phosphoproteome (including TiO_2_ enriched phospho-serine / threonine and anti-phophotyrosine antibody-enriched phosphotyrosine peptides) analyses were performed. Overall, we identified and quantified thousands of proteins and phosphosites (**Fig. 1B and Table S1, S2**). Approximately 66-69% of the phosphosites identified were Class I sites. These are phosphosites whose localization probability of phosphorylation is greater than or equal to 0.75. More proteins and phosphosites were identified from the H1975/AZR3/AZR4 experiment than the H1975/COR1/COR10 experiment. 77% of the proteins and 39% of the phosphosites identified were common to both the osimertinib and rociletinib experiments. More unique proteins and phosphosites were identified in the osimertinib experiment than the rociletinib experiment (**Fig. 1C**). This could be due to the use of the QE HF mass spectrometer for the osimertinib experiments, compared to the slower, less sensitive orbitrap elite for the rociletinib experiments. In addition, more biological replicates were performed for the osimertinib experiments, especially for the phosphotyrosine enrichment. The hierarchical clustering of proteins and phosphosites (**Fig. 1D**) clearly showed that the two drug resistant cell lines (osimertinib and rociletinib) were grouped into two separate clusters. The osimertinib resistant cell lines (AZR3 and AZR4), with or without drug treatment, were clustered together as were the rociletinib resistant cell lines (COR1 and COR10). Overall, more proteins and phosphopeptides were less abundant in the resistant cells as evidenced by the distribution of the log_2_ ratio of the resistant and sensitive cells (**Figure S1**) and the actual numbers of significantly altered proteins and phosphopeptides (**Figure S2A**). The two rociletinib resistant cell lines, COR1 and COR10 were similar to each other and there was greater difference between AZR3 and AZR4, the two osimertinib resistant cells (**Figure S2A**).

**Fig. 1.**
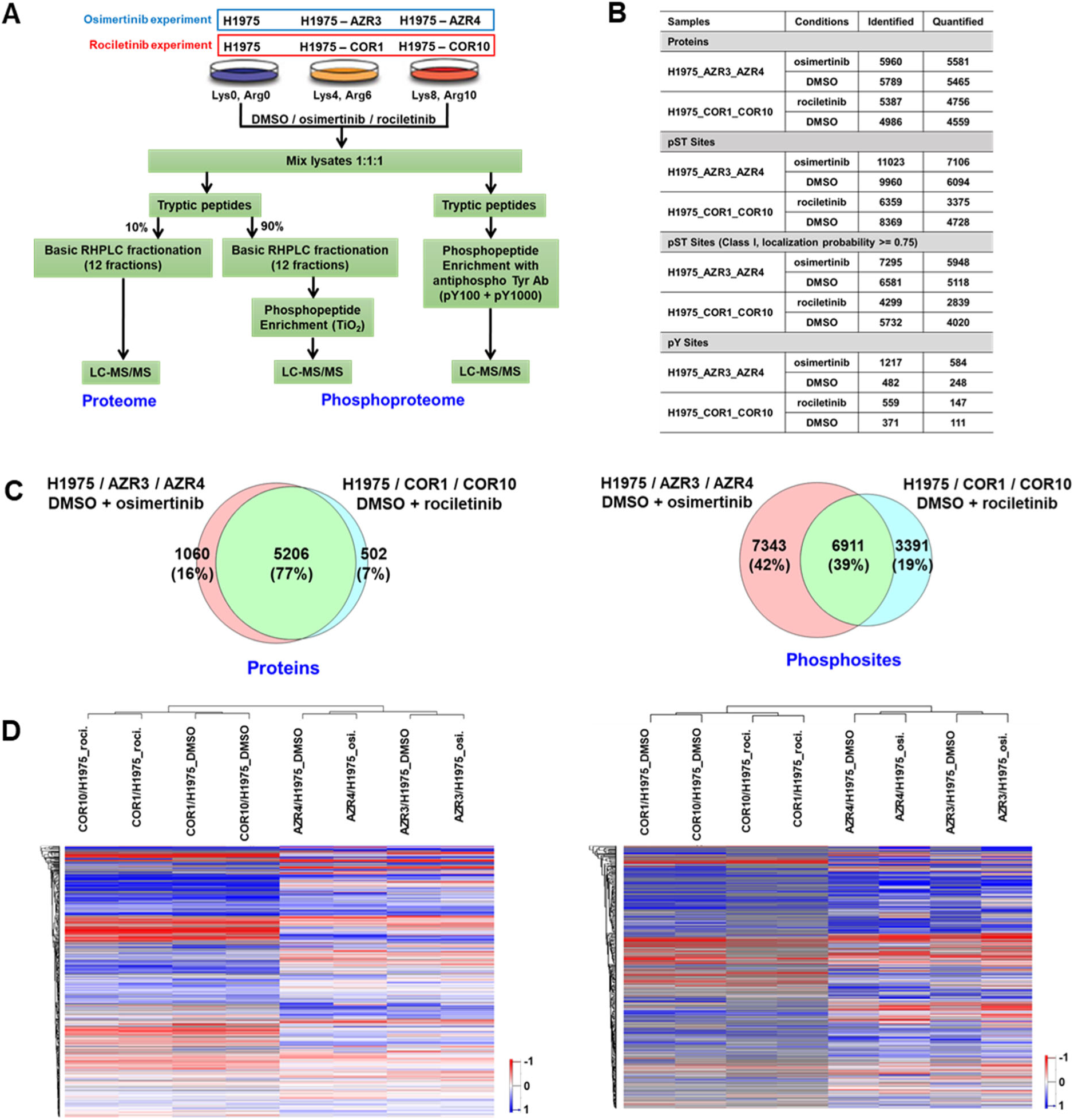
Summary of SILAC-based quantitative proteome and phosphoproteome analyses of isogenic 3^rd^ generation EGFR TKI-sensitive and resistant lung adenocarcinoma cells. (A) Experimental workflow showing treatment of SILAC-labelled cells, enrichment of phosphopeptides, and detection by tandem mass spectrometry. TKI-sensitive H1975, osimertinib-resistant AZR3/AZR4 cells, and rociletinib-resistant COR1/COR10 cells were treated with DMSO or the corresponding TKI. (B) Summary table showing the number of proteins and phosphosites identified in each experiment. (C) Venn diagrams of proteins (left panel) and phosphosites (right panel) identified in osimertinib and rociletinib experiments. (D) Hierarchical clustering of proteins (left panel) and phosphosites (right panel) based on SILAC ratios. Columns represent different cell lines treated as indicated. Rows represent quantified proteins or phosphopeptides identified in all experimental conditions.

Next, one sample t-tests were performed on the protein and phosphopeptide SILAC ratios to determine significant differences in abundance between the resistant and sensitive cells. Hundreds of proteins and phosphosites were identified whose abundances wer significantly different in resistant and sensitive cell lines (P < 0.05, 1.5-fold change) (**Figure S2A-B, Table S2**). We generated volcano plots using the log_10_ (p value) and log_2_ (fold change) of the proteins and phosphosites (**Figure S3**). The abundance of 113 proteins and 79 phosphosites were significantly altered in osimertinib resistant AZR3 and AZR4 cells treated with either DMSO or osimertinib. 463 proteins and 122 phosphosites were significantly altered in rociletinib resistant cells, COR1 and COR10, treated with either DMSO or rociletinib. Among the proteins differentially expressed proteins in the resistant cell lines were kinases, phosphatases, transcription regulators, transporters, and enzymes (**Figure S2B**). Examination of protein localization and function for all significantly altered proteins by Gene Ontology (GO) classification analyses showed that the differentially regulated proteins localized to the cytoplasm, nucleus, plasma membrane and the extracellular space (**Figure 2A**). Many translation regulator proteins were significantly more altered in rociletinib compared to osimertinib-resistant cells (**Table S3**). The heatmap of selected translation regulators illustrates the altered expression of proteins including several EIF proteins (**Figure 2B**).

**Fig. 2.**
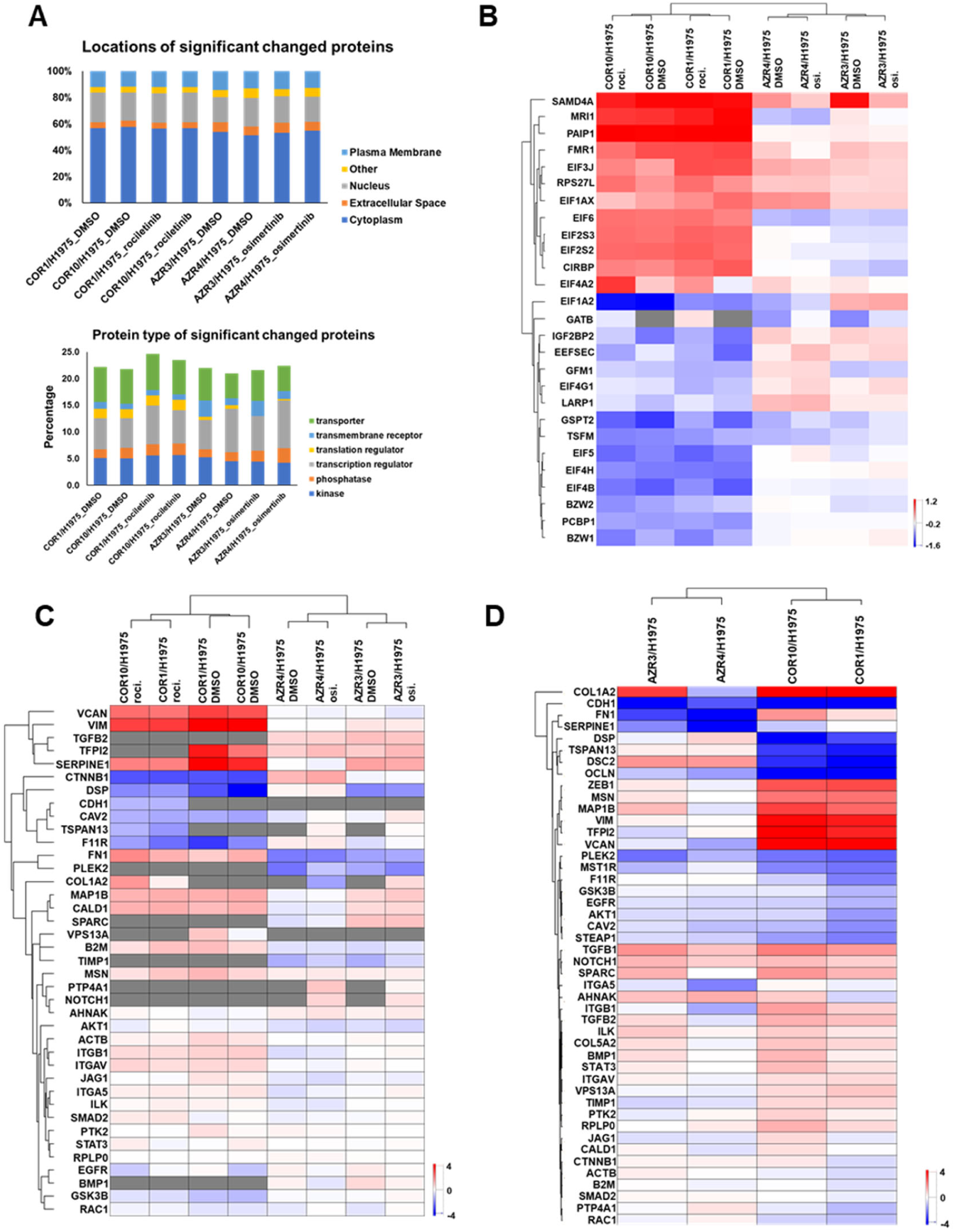
GO classification analyses for localization and function of proteins with altered abundance, translation regulators, EMT proteins and transcripts in TKI-resistant cells. (A) Percentage of differentially expressed proteins in TKI-resistant cells with specific subcellular localization (up) and function (bottom). (B) Heatmap of SILAC ratios of protein abundance (TKI-resistant/sensitive) of selected translation regulators across all experiments demonstrate significantly more alterations in rociletinib-resistant cells. (C) Hierarchical clustering of quantified EMT proteins in three state SILAC experiments based on SILAC ratios of protein abundance in presence and absence of corresponding TKI. (D) Hierarchical clustering of transcript ratios of EMT genes quantified by qPCR.

Epithelial-mesenchymal transition (EMT) is a biological program observed in several types of epithelial cancers including NSCLC. It has been associated with metastatic spread and EGFR inhibitor resistance (*20-22*). We asked whether the osimertinib and rociletinib resistant cells underwent EMT and whether we can identify an EMT protein signature from the mass spectrometry data. We evaluated the expression of EMT proteins across all the resistant cell lines. In addition, we examined the expression of a set of EMT transcripts by quantitative RT-PCR (**Table S4**). At both the protein and RNA level, the ratio of TKI-resistant/sensitive cells in the presence and absence of the corresponding TKI (**Figure 2C-D**) showed that the expression of many EMT signature genes, such as vimentin (VIM) and Fibronectin 1 (FN1), was modulated consistent with EMT in resistant cells. This pattern of EMT genes expression was more significant in rociletinib-resistant cells. E-cadherin protein (CDH1) expression, associated with the epithelial state, has been associated with longer time to progression and a trend toward longer overall survival following combination erlotinib/chemotherapy (*23*). Consistent with this, we observed lower protein and transcript expression of CDH1 in resistant cells. Our results suggest that EMT is associated with TKI drug resistance in both rociletinib and osimertinib-resistant cells.

### Changes in the abundance and phosphorylation of protein kinases and phosphatases in osimertinib and rociletinib resistant cells

To further examine the altered protein expression of kinases and phosphatases identified (**Figure 2A**), the significantly altered SILAC ratios for kinases (**Figure S4A)** and phosphatases (**Figure S4B)** in the resistant cell lines, both DMSO and drug treated, were visualized in a heatmap. The significantly altered proteins (p<0.05) with a fold change cut-off of 1.5 were chosen. There were many kinases and phosphatases with altered expression in the TKI-resistant cells. There were more proteins dysregulated in the rociletinib resistant cells than osimertinib resistant cells, especially those expressed at lower levels. Expression of protein tyrosine kinase 7 (PTK7), an inactive tyrosine kinase involved in Wnt signaling (*24-27*), and two phosphatases, translocase of inner mitochondrial membrane 50 (TIMM50) and protein tyrosine phosphatase receptor type F (PTPRF), was increased in all resistant lines. Abundance of several kinases, such as thymidine kinase 1 (TK1), P21 activated kinase 1 (PAK1), phosphatidylinositol-5-phosphate 4-kinase type 2 gamma (PIP4K2C), and phosphatases, RNA guanylyltransferase and 5’-phosphatase (RNGTT), protein phosphatase, Mg^2+^/Mn^2+^ dependent 1B (PPM1B) and protein phosphatase 1 regulatory inhibitor subunit 14B (PPP1R14B) was decreased in all resistant lines. Interestingly, several proteins displayed opposite patterns in the TKI-resistant cell lines, such as 5’-Nucleotidase Ecto (NT5E), was more abundant in osimertinib-resistant, but less abundant in rociletinib-resistant cells, in comparison to their sensitive counterparts.

There were also differences in expression of certain kinases and phosphatases between the two different isogenic cell lines resistant to either osimertinib (AZR3 and 4) and rociletinib (COR1 and 10). For example, we saw contrasting expression patterns for the SRC proto-oncogene, an important non-receptor tyrosine kinase activated downstream of RTK, which plays a role in the activation of other protein tyrosine kinase (PTK) families. Whereas expression of SRC was increased in both the COR1 and COR10 rociletinib resistant cells and the osimertinib resistant cell line AZR3. but decrease in the resistant cell line AZR4. Such difference in individual protein expression in isogenic TKI-resistant cells, such as AZR3 and 4 may be represent their differential modulation during resistance. The phosphatase, Inositol Polyphosphate-5-Phosphatase (OCRL) involved in regulating membrane trafficking was less abundant in the AZR4 cells only. N-Myc Downstream Regulated 1 (NDRG1) had lower expression in the resistant cell lines, AZR4 and COR1. Overall, the differences in the modulation of the expression of specific kinases and phosphatases among isogenic versions of TKI-resistant cells underscore the heterogeneity in the development of resistance to the EGFR TKIs, osimertinib and rociletinib.

Since phosphorylation is regulated by both kinases and phosphatases, which are themselves regulated by phosphorylation, we evaluated the altered phosphorylation of specific phosphosites on kinases and phosphatases in the resistant cells. Heatmaps of the SILAC phosphorylation ratios (resistant/sensitive) highlight significant alterations in the phosphorylation of individual sites in kinases (**Figure S5A)** and phosphatases **(Figure S5B)** across the TKI-resistant cell lines with/without TKI treatment. Phosphorylation of many important sites on kinases, including EGFR-Y1172, MET-Y1003/Y1234, GAB1-Y659/Y406, MAPK1-Y187 was significantly reduced in resistant cells, independent of TKI treatment. On the other hand, phosphorylation of many sites, such as CDK1-Y15, CDK2-T14/Y15, CHEK2-S379, ROCK2-S1374, was significantly increased in resistant cells. Interestingly, we identified phosphorylation sites within several kinases which were differentially modulated with and without TKI treatment. In osimertinib-resistant cells, reduced phosphorylation of MAPK3K2-S154, EGFR-Y1172, HIPK-Y359, YES1-Y194, EPHB4-Y590, CDK5-Y4, GAB1-Y405/Y659, LYN-Y32 was more pronounced in untreated as compared to osimertinib-treated resistant cells. Similarly, in rociletinib-resistant cells, reduced phosphorylation of MARK3-S469, TJP2-S986, MAST4-S1373, SRP72-S621, MTOR-T1162, BAIAP2-T360, TNK2-Y859, STK10-S450, AAK1-S624, compared to rociletinib treated resistant cells. These dynamic phosphorylation changes suggest that, in TKI-resistant cells, phosphorylation at these sites is less dependent on EGFR signaling.

We also investigated changes in phosphorylation on phosphatases (**Figure S5B**). The phosphatase, PTPN11, located downstream of EGFR, regulates the RAS/MAPK signaling pathway. Phosphorylation of PTPN11-Y62 was reduced in both osimertinib and rociletinib resistant cells, regardless of TKI treatment. In rociletinib resistant cells phosphorylation of PTPN11-Y584 was also reduced. We also identified another differentially modulated phosphatase, PTPN3. Interestingly, PTPN3 has been shown to be upregulated in both cisplatin and doxorubicin-resistant ovarian cancer cells, suggesting its role in tumorigenesis, stemness and drug resistance in ovarian cancer and potential its use as a therapeutic target for ovarian cancer (*28*). Phosphorylation of PTPN3-S359 was significantly reduced in rociletinib resistant cells, COR1 and COR10 but increased in osimertinib resistant cells, AZR3 and AZR4. Integrin-linked Kinase-Associated Serine Threonine Phosphatase (ILKAP) also known as protein phosphatase 2C selectively associates with Integrin Linked kinase (ILK) and modulates cell adhesion and growth factor signaling (*29*). ILKAP inhibits S9 phosphorylation of GSK3β. ILKAP-S13 phosphorylation was reduced significantly in osimertinib resistant cells, AZR3 and AZR4. Taken together, we identified altered phosphorylation of key phophosites on kinases and phosphatases implying that resistance is accompanied by dysregulation of kinase and phosphatase activity and in turn the relevant signaling pathways.

### Protein abundance differences between osimertinib and rociletinib-resistant cells in presence of the respective TKI

Osimertinib and rociletinib are both 3rd generation EGFR TKIs with similar mechanisms of action. They are both irreversible inhibitors which covalently bind C797 of mutant EGFR, and do not inhibit wild type EGFRs. However, small molecule inhibitors such as TKIs often have different off target effects that have implications for targeted therapy and resistance. We sought to specifically analyze the proteins whose modulation is similar or different upon resistance to both TKIs. We examined proteins with significantly altered expression in all four TKI-resistant cell lines; the osimertinib resistant cell lines AZR3 and AZR4 as well as the rociletinib resistant cell lines, COR1 and COR10. There were 60 proteins with altered expression in all four TKI resistant cell lines treated with their corresponding TKIs, osimertinib and rociletinib (**Figure 3A and Table S5**). The heatmap and hierarchical clustering of the 60 common differentially expressed proteins shows that the protein expression is either increased or decreased in both COR1 and COR10 cells or AZR3 and AZR4 cells, suggesting that these proteins are likely involved in the resistance mechanisms to the individual EGFR TKIs. Of these 60 proteins, 35 were less abundant in all resistant lines, while nine proteins were expressed at significantly higher levels in all resistant lines. Interestingly, expression of 16 proteins were differentially modulated in the osimertinib and rociletinib resistant cells. Of these proteins, expression of 15 increased in rociletinib resistant cells and decreased in osimertinib resistant cells compared to the isogenic TKI sensitive cells (**Figure 3B**). Included in these group is fibronectin 1 (FN1), Transglutaminase 2 (TGM2), Complement C3 (C3), HLA-A and Inosine Triphosphatase (ITPA). Only one protein, ALCAM, a member of a subfamily of immunoglobulin receptors, was down-regulated in rociletinib resistant cells but up-regulated in osimertinib resistant cells. STRING analysis of the 60 common differentially expressed proteins revealed a network of proteins in which 27 of the 60 are direct or indirect downstream targets of EGFR (**Figure 3C**), indicating EGFR downstream targets are alterd in the EGFR TKI-resistant cells.

**Fig. 3.**
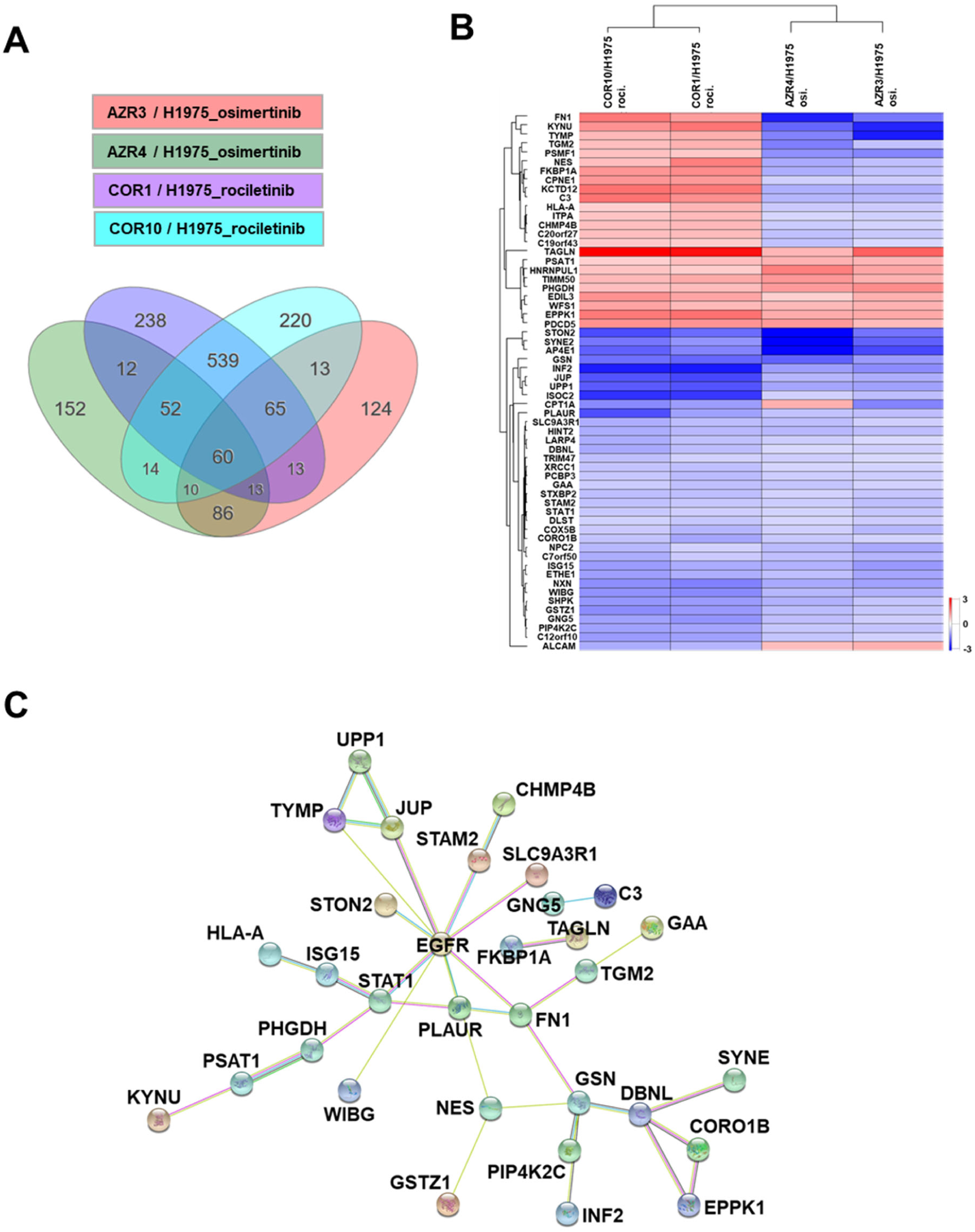
Comparison of protein abundance differences between osimertinib-resistant and rociletinib-resistant cells treated with respective TKI. (A) Differentially expressed proteins in TKI-resistant cells compared to the sensitive cells. 60 differentially expressed proteins were common to all four resistant cell lines. (B) Heatmap and hierarchical clustering of the 60 differentially expressed proteins common to all TKI-resistant cell lines. (C) Protein network of the 60 common differentially expressed proteins together with EGFR by STRING analysis. Many of these proteins are direct and indirect downstream targets of EGFR.

### Canonical pathways dysregulated in osimertinib and rociletinib resistant cells

Next, we used Ingenuity Pathway Analysis (IPA) to evaluate the activated or inhibited canonical pathways enriched among the proteins with significantly altered phosphorylation (**Table S6**). Interestingly, a number of signaling pathways enriched in both the osimertinib resistant cells (**Figure S6A**) and rociletinib resistant cells (**Figure S6B**) had a negative Z-score, which is indicative of pathway inhibition. In both TKI-resistant lines, ERBB signaling, EGF signaling, regulation of eIF4 and p70S6K signaling, ILK signaling, JAK/STAT signaling, RAC signaling, and PAK signaling were inhibited. Many of the pathways enriched in the resistant cells had positive Z-scores, suggesting activation, including PTEN, PPARγ, RHOA, RhoGDI, p53, and PPARα/RXRα signaling. There were a few signaling pathways, in the osimertinib and rociletinib resistant cells, which were dysregulated in an opposing manner. For example, CDC42 signaling and regulation of Actin-based motility by RHO pathway signaling were activated in the osimertinib resistant cells and inhibited in the rociletinib resistant cells. Overall, we found that, more signaling pathways were inhibited in the resistant cells, correlating with the overall significant reduction of phosphorylation in the resistant cells.

### Validation of differentially phosphorylated proteins and phosphosites in resistant cells

To further validate and supplement our global quantitative mass spectrometry data, we performed Western blots of select proteins (**Figure 4**). Changes in protein expression and phosphorylation at specific phosphosites in TKI-sensitive cells, H1975 and the TKI-resistant cells, COR1, COR10, AZR3, and AZR4 with or without TKI treatment, were examined (**Figure 4A)**. Relative quantification of phosphorylated proteins normalized to total protein expression was performed on the Western blots. Phosphorylation of EGFR, ERK, AKT-T308, SRC-Y416, MEK-S217/221 and p70S6K-T389 was reduced in at least one of the resistant cell lines. Phosphorylation of MET, an RTK with known activation of parallel signaling pathway upon EGFR TKI resistance, was increased in COR10 and both AZR3 and AZR4. AXL expression was upregulated in both the osimertinib and rociletinib resistant cells; however, AXL-Y779 phosphorylation was reduced in AZR3 and AZR4 cells without osimertinib treatment, but not significantly in COR1 and COR10 cells. Among the epithelial-mesenchymal transition (EMT) proteins, E-cadherin expression consistently decreased in both the osimertinib and rociletinib resistant cell lines, again confirming EMT associated with the resistant cells (**Figure 4A-B**).

**Fig. 4.**
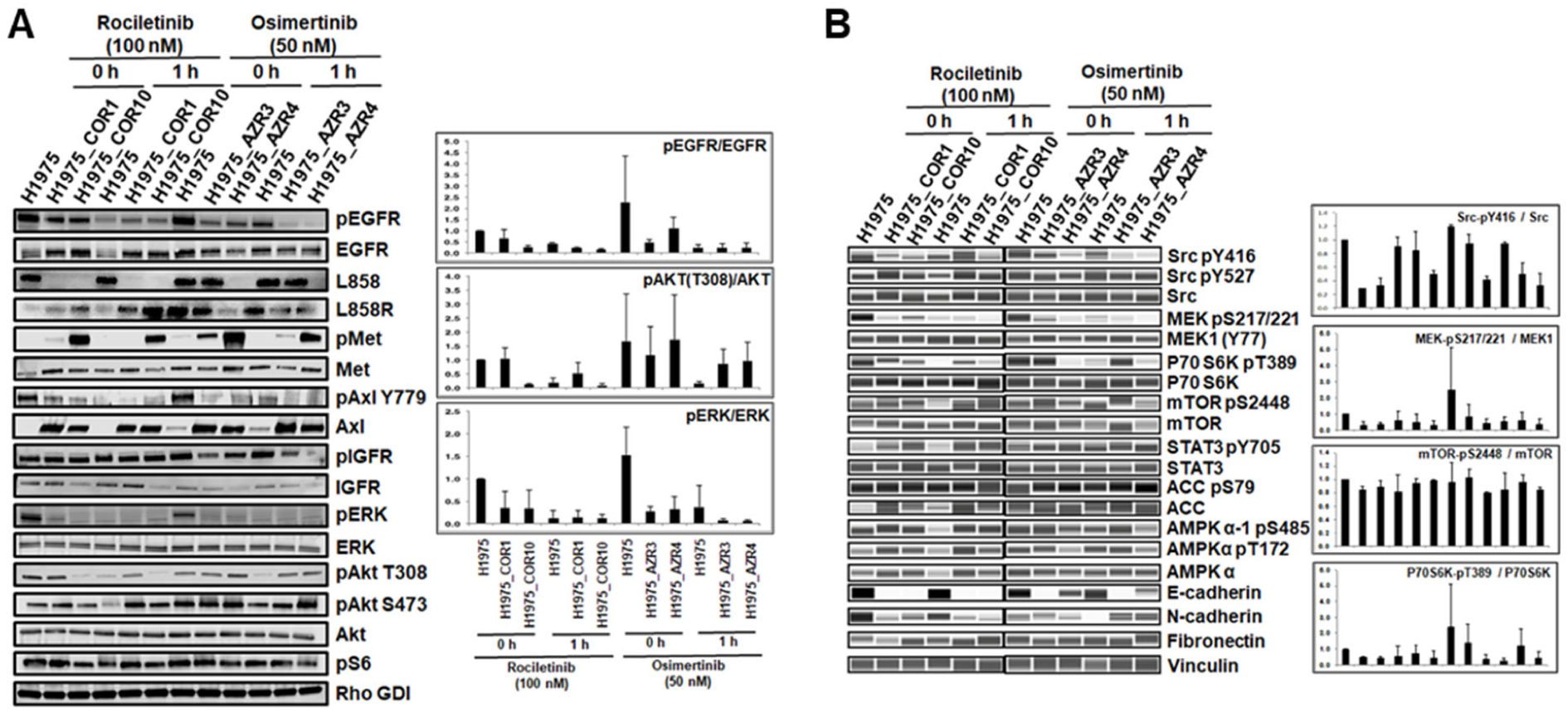
Validation of differentially phosphorylated phosphosites and differentially expressed proteins in osimertinib or rociletinib-resistant cells. (A-B) Western blots showing changes in phosphorylation and total protein expression without TKI treatment and upon 1 hour of rociletinib (100 nM) or osimertinib (50 nM) treatment in H1975, COR1, COR10, AZR3, and AZR4 cells. Bar graphs show the relative quantification of phosphorylated proteins normalized to total protein expression in each cell line.

### Upstream phosphatase analysis of significantly altered phosphopeptide substrates in TKI-resistant cells

Phosphorylation of downstream targets is regulated by upstream kinases and phosphatases. To identify the upstream regulatory phosphatases, we performed IPA analysis on the phosphorylation sites that were significantly hyper- or hypo-phosphorylated in the resistant cells. We identified 13 osimertinib-specific (Figure S7A) and 14 rociletinib-specific **Figure S7B**) upstream phosphatases in the resistant cells. Almost all had positive Z-scores, indicating they are active in the TKI-resistant cell lines.

Next, we made networks showing connections between altered phosphosites and their experimentally validated phosphatases from the human DEPhOsphorylation Database (DEPOD) (*30*) (**Figure S7C-F**). Many of these phosphatases, such as PTPN11, PTPN1, PTPRK, and PTPRJ, overlapped with those identified by IPA analysis. Phosphorylation of downstream phosphorylation targets of these phosphatases was reduced in the resistant cells, suggesting activation of these phosphatases.

### Validation of altered phosphorylation sites of the phosphatase PTPN11

The phosphatase, PTPN11 (more commonly known as SHP2) can activate the RAS/MAPK pathway and inhibit the PI3K/AKT cascade through different mechanisms. Once SHP2 is associated with adaptor protein GAB1, it inhibits PI3K by dephosphorylating GAB1-PI3K binding sites. We asked whether phosphorylation changes in SHP2 are associated with PI3K/AKT and MAPK signaling in TKI resistant cells. Multiple phosphorylation sites of PTPN11 were identified and quantified in both osimertinib and rociletinib resistant cells (**Figure 5A**). Most of the sites identified, including S580, Y546, Y584, and Y62, were significantly hypo-phosphorylated in all resistant cells,. MS and MS/MS spectra of the PTPN11 phosphopeptides VpYENVGLMQQQK (Y584) and IQNTGDpYYDLYGGEK (Y62) demonstrate that phosphorylation at these sites was significantly lower in all resistant cells, AZR3 and AZR4 as well as COR1 and COR10 (**Figure 5B**). Western blots confirmed the PTPN11phosphorylation changes in sensitive and resistant cells with or without drug treatment. PTPN11-Y584 phosphorylation was significantly reduced in both the osimertinib and rociletinib resistant cells (**Figure 5C**). Taken together, reduced phosphorylation of key PTPN11sites in resistant cells, which is consistent with inactive phosphatase activity, is associated with increased activation of PI3K/AKT pathway and decreased activation of the RAS/MAPK signaling (**Figure 5D**).

**Fig. 5.**
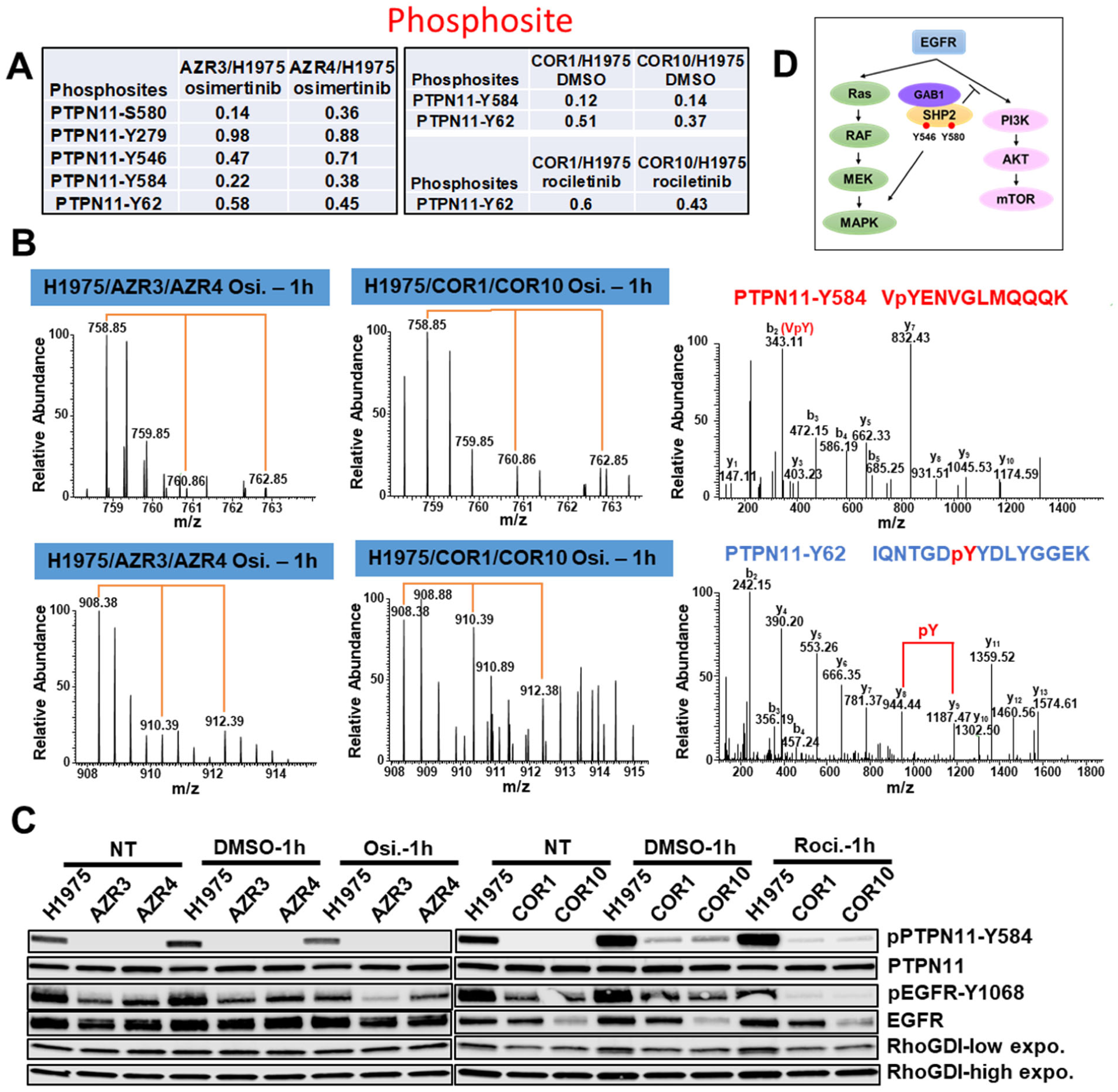
Phosphorylation alteration of different phosphosites of the phosphatases PTPN11. (A) Multiple phosphorylation sites of PTPN11 identified and quantified in both osimertinib and rociletinib resistant cells. The ratios give the relative abundance between the resistant cells and the sensitive cells H1975. (B) Annotated MS/MS spectra of the phosphopeptides VpYENVGLMQQQK (Y584) and IQNTGDpYYDLYGGEK (Y62) of PTPN11 and the MS spectra of their parent ions showing the relative abundance changes between the sensitive and resistant cells in osimertinib and rociletinib experiments. (C) Western blots showing changes in phosphorylation and total protein expression of PTPN11 and EGFR without TKI treatment and upon 1 hour of rociletinib (100 nM) or osimertinib (50 nM) treatment in H1975, COR1, COR10, AZR3, and AZR4 cells. (D) Role of SHP2 in the activation of RAS/MAPK and PI3K/AKT signaling pathways downstream from EGFR.

### Upstream kinase analysis of significantly altered phosphopeptide substrates in TKI-resistant cells

To identify kinases whose targets were significantly over-represented among the significantly hyper- or hypo-phosphorylated sites in drug-resistant vs. sensitive cells, we matched the significantly altered phosphorylation sites with the kinase-target site data in iPTMnet v.4.1 (*31*). To improve statistical power, the enrichment analysis was performed at the level of kinase families rather than individual kinases (**Tables S7, S8**). Networks were generated showing the upstream kinase families and the downstream phosphorylated targets that were either significantly hyper- or hypo-phosphorylated in AZR3, AZR4, COR1 and COR10 cell lines in presence of the corresponding EGFR TKIs (**Figure 6**). We determined that many downstream CDK kinase family phosphosites in the CMGC group were hypo-phosphorylated; while many downstream AKT kinase family and STE20 kinase family phosphosites were hyper-phosphorylated in all four resistant cell lines (**Figure 6A-D**). We also found phosphosites in proteins downstream of the PKA kinase family were hyper-phosphorylated in the two rociletinib resistant cell lines (**Figure 6C, D**). There was more similarity between networks identified for the rociletinib-resistant lines, COR1 vs COR10, than between the osimertinib resistant cell lines, AZR3 vs AZR4. The CK2, SGK and WEE kinase families downstream proteins were hyper-phosphorylated in the AZR3 resistant cell line, (**Figure 6A**). Phosphosites downstream of several CAMK group (CAMK1, RAD53, and MAPKAPK) and the TK group (ACK and MET) kinase families were either hyper- or hypo-phosphorylated in the other osimertinib resistant cell, AZR4 (**Figure 6B**). In summary, global quantitative phosphoproteomic analyses have identified key regulatory networks consisting of the upstream kinase families and the downstream phosphorylation targets that are potentially activated or inhibited in EGFR TKI resistant cells.

**Fig. 6.**
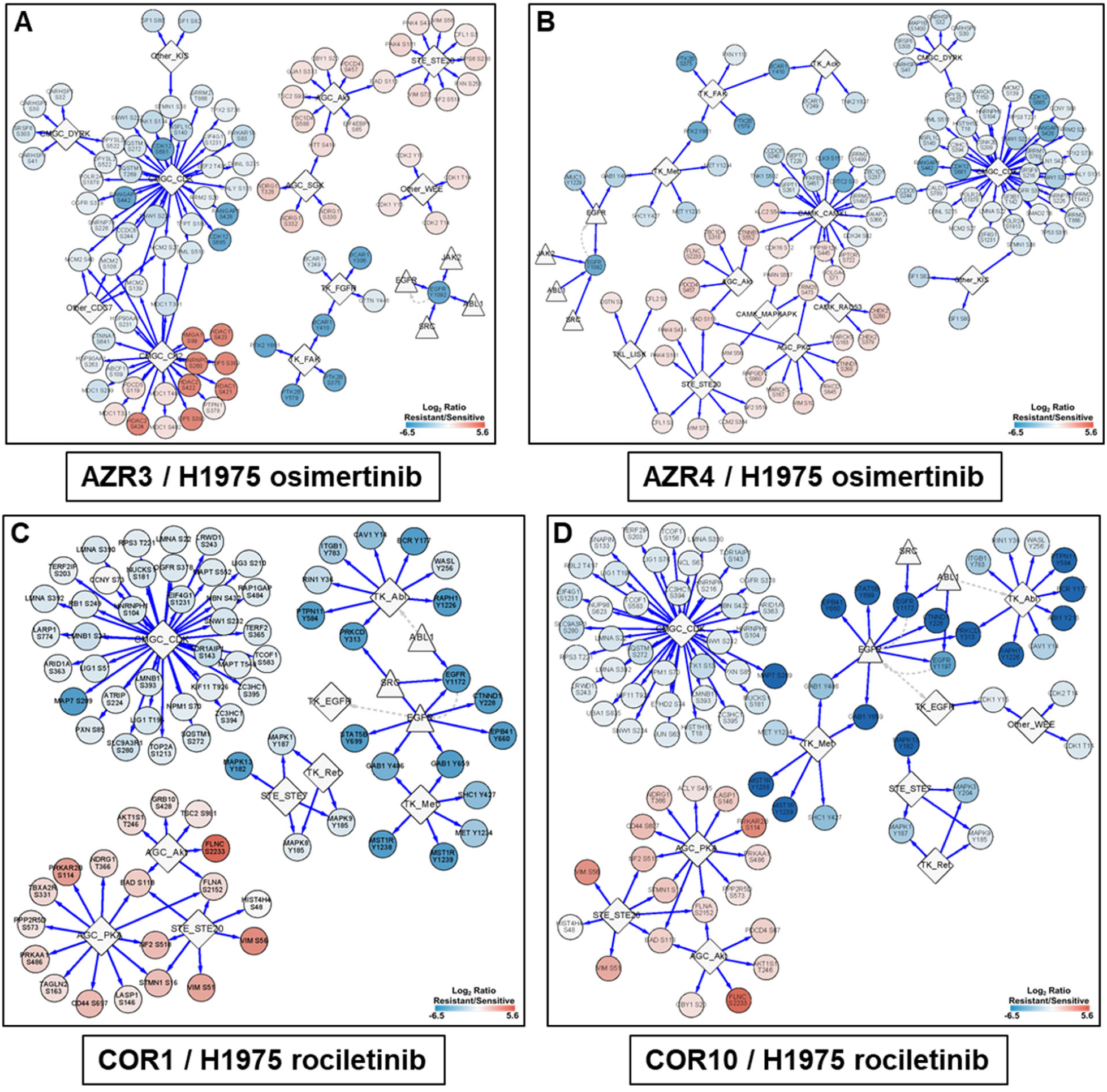
Networks generated by upstream kinase analysis of significantly altered phosphopeptide substrates in TKI-resistant cells using iPTMNet. (A) AZR3/H1975 in the presence of osimertinib; (B) AZR4/H1975 in the presence of osimertinib; (C) COR1/H1975 in the presence of rociletinib; (D) COR10/H1975 in the presence of rociletinib. The scale bars show the log_2_ SILAC ratio of phosphorylation (resistant/sensitive). Phosphosites in red are hyperphosphorylated and the ones in blue are hypophosphorylated in the resistant cells.

### Prediction of drugs to circumvent resistance and in vitro / in vivo efficacy testing in osimertinib resistant cells

Next, we used the P100 phosphoproteomic data generated from cancer cell lines with and without drug treatment by the NIH Library of Integrated Network-Based Cellular Signatures (LINCS) program (http://www.lincsproject.org/) (*32*) to identify drugs that might reverse the phosphoproteomic signature from the TKI resistant cells. We looked for anti-correlation of the phosphoproteomic signatures from osimertinib and rociletinib resistant cells with specific P100 phosphoproteomic signatures from cell lines treated with specific drugs. We postulated that the drugs whose phosphorylation signatures are anticorrelated with those from the TKI resistant cell lines will have the potential to circumvent resistance to the EGFR TKIs, osimertinib and rociletinib. Using this approach, several inhibitors of multiple signaling pathways were identified for both osimertinib and rociletinib resistant cells (**Table S9**). We selected dactolisib, a dual PI3K/AKT and MTOR inhibitor and VX-970, a DNA repair pathway inhibitor for further validation. Dactolisib could overcome TKI resistance in both osimertinib and rociletinib resistant cells either by itself or in combination with rociletinib or osimertinib (**Figure 7A**). VX-970 was also found to have efficacy in circumventing TKI resistance in osimertinib resistant cells, AZR3 and AZR4 (**Figure 7B**). We further investigated whether dactolisib and VX-970 could inhibit xenograft tumor growth *in vivo* in mice. Mice were injected with AZR3 cells to develop tumors in flank. Dactolisib in combination with osimertinib was able to inhibit tumor growth of AZR3 xenografts *in vivo* (**Figure 7C**) while VX-970 in combination with osimertinib could not inhibit tumor growth *in vivo* (data not shown). The results show the feasibility of predicting drugs that overcome TKI resistance by leveraging the phosphoproteomic signature of drug treatment in cancer cells from the LINCS program.

**Fig. 7.**
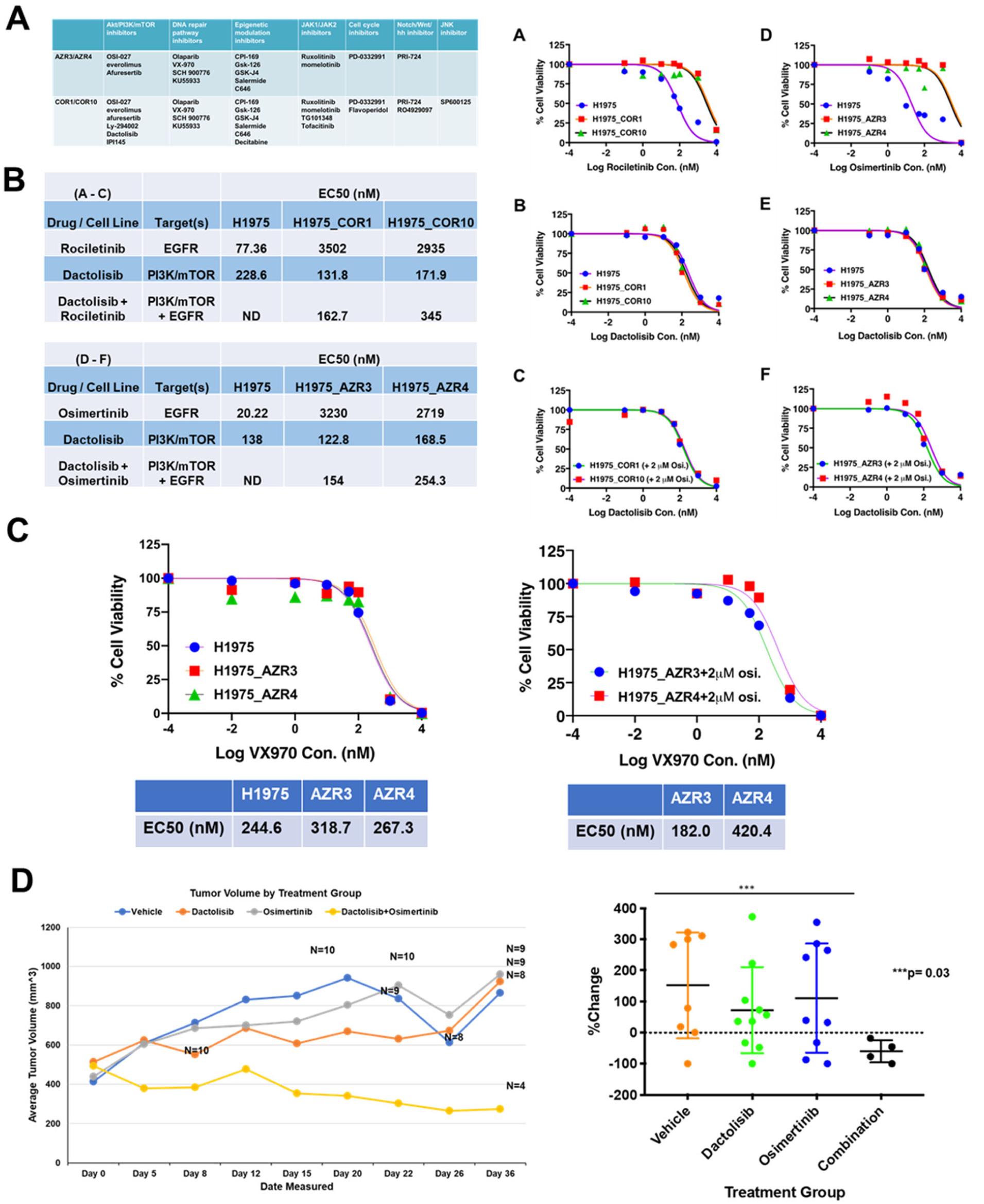
*In vitro* and *in vivo* sensitivity of osimertinib-resistant cells to dactolisib, a PI3K/mTOR inhibitor and VX970, an ATR inhibitor. (A) Drugs predicted to overcome the drug resistance by NIH LINCS Program. (B) EC_50_ of dactolisib shows the efficacy of dactolisib in circumventing osimertinib resistance. (C) EC_50_ of VX970 shows the efficacy of VX970 in circumventing TKI resistance. (D) Dactolisib in combination with osimertinib inhibits tumor growth of H1975-AZR3 xenografts *in vivo*.

## Discussion

The use of oral EGFR TKIs for the treatment of lung cancer patients with tumors harboring mutant EGFR has been a paradigm shift. Osimertinib, the 3^rd^ generation EGFR TKI is the current standard of care for patients with EGFR mutations due to its increased efficacy, lower side effects, and increased brain penetrance. Unfortunately, all patients treated with this drug develop resistance. Genomic approaches have primarily been used to interrogate resistance mechanisms to EGFR TKIs, including the 3^rd^ generation EGFR TKIs. Here, we have characterized the proteome and the phosphoproteome of a series of isogenic EGFR mutant lung adenocarcinoma cell lines that are either sensitive or resistant to the 3^rd^ generation EGFR TKIs, osimertinib and rociletinib. To our knowledge, this is the most comprehensive proteomic dataset resource to date used to investigate the 3^rd^ generation EGFR TKI resistance in lung adenocarcinoma. We have utilized this unbiased global quantitative proteomic and phosphoproteomic dataset to uncover alterations in signaling pathways in the TKI-resistant lines. In addition, we used this resource to reveal a proteomic EMT signature as well as kinases and phosphatases with altered protein expression and phosphorylation in the TKI resistant cells. Furthermore, we used anticorrelation analysis of this phosphoproteomic dataset with the published drug-induced P100 phosphoproteomic datasets from the LINCS analysis to postulate drugs with potential to overcome EGFR TKI resistance. We validated one such drug, dactolisib, a PI3K/mTOR inhibitor, in combination with osimertinib was able to overcome osimertinib resistance both *in vitro* and *in vivo*.

Osimertinib and rociletinib resistant cells were independently developed from H1975 cells which harbor the EGFR-L858R/T790M mutation, by stepwise increases in drug dosage over several months (*3, 33*). The sensitive cells phenocopy approximately 60% of patients who develop resistance to the 1^st^ and 2^nd^ generation EGFR TKIs via acquisition of the EGFR-T790M mutation. Furthmore, the resistant cells serve as a valid model system for patients who undergo osimertinib treatment, and eventually develop resistance. We and others have primarily used genomic strategies to investigate mechanisms of osimertinib resistance (*9, 11-15*). To our knowledge, this is the first study to identify and compare resistance mechanism of these 3^rd^ generation EGFR TKIs by utilizing unbiased global proteome and phosphoproteome modification data. This approach provides a powerful tool to examine activated signaling pathways without bias.

The experimental approach used in this study employed 3-6 global SILAC quantitative mass spectrometry biological replicates. We identified thousands of proteins and phosphosites in both experiments, hundreds of which had statistically significant alterations in the resistant cells. Surprisingly, we observed more proteins and phosphosites with decreased abundance in both osimertinib and rociletinib resistant cells with/without drug treatment. Moreover, many translation regulator proteins were significantly more altered in rociletinib-resistant cells compared to osimertinib-resistant cells, such as several of the EIF proteins (EIF3J, EIF1AX, EIF6, EIF2S3, EIF2S2, EIF4A2, EIF1A2, EIF4G1, EIF5, EIF4H and EIF4B), RPS27L, EEFSEC, IGF2BP2, PAIP1 (**Figure 2B**). The results suggest that the regulation of translation may have been altered in rociletinib resistant cells in contrast to the osimertinib resistant cells. Global alteration of protein translation would result in changes in protein expression contributing to TKI resistance. Furthermore, translation could be used as a therapeutic target (*34*).

EMT has been associated with resistance to targeted therapies, including EGFR TKIs (*20, 21*). A transcriptomic EMT signature has been validated in clinical studies (*21*). We investigated quantitative transcript expression of EMT genes using qPCR and further estimated the protein expression changes in TKI resistant cells from the quantitative mass spectrometry data. Many EMT signature proteins and transcripts were altered significantly in the resistant cells, especially in the rociletinib resistant cells. For example, vimentin (VIM) and fibronectin 1 (FN1) were more abundant and E-cadherin protein (CDH1) expression was reduced in TKI-resistant cells. Our results suggest that EMT is associated with TKI drug resistance for both rociletinib and osimertinib-resistant cells. We propose an EMT protein signature that can be identified and analyzed using mass spectrometry. This EMT protein signature needs to be further validated with large datasets employing targeted mass spectrometry assays, such as multiple reaction monitoring (MRM) in a QQQ mass spectrometer (*35-37*).

We observed more differences between the two isogenic osimertinib resistant cells, AZR3 and AZR4 compared to the two rociletinib resistance cells, COR1 and COR10. The iPTMnet of upstream kinase analysis showed that downstream protein targets of the CK2, SGK and WEE kinase families were hyper-phosphorylated in the resistant cell line, AZR3 (**Figure 6A**). Phosphosites downstream of several kinase families in the CAMK group (CAMK1, RAD53, and MAPKAPK) and the TK group (ACK and MET) were either hyper- or hypo-phosphorylated in the other osimertinib resistant cell, AZR4 (**Figure 6B**). To the contrary, the two rociletinib resistant cells, COR1 and COR10 were more similar to each other with respect to the upstream kinases and the downstream targets. These results obtained from isogenic TKI-resistant cells underscore the proteomic heterogeneity that is likely to impact treatment response and resistance.

We have identified and quantified many kinases and phosphatases with altered expression and phosphorylation in the osimertinib and rociletinib resistant cells. Moreover, many important phosphorylation sites on kinases were hyper- or hypo-phosphorylated significantly in the resistant cells, both with and without drug treatment. Many of them are important downstream EGFR signaling pathway proteins, including EGFR-Y1172, MET-Y1003/Y1234, GAB1-Y659/Y406, MAPK1-Y187, CDK1-Y15, CDK2-T14/Y15, CHEK2-S379, ROCK2-S1374.

PTPN11, commonly known as SHP2, is a ubiquitously expressed well characterized PTP oncogene. SHP2 activates the RAS/MAPK signaling pathway and inhibits the PI3K/AKT cascade downstream of RTKs by various mechanisms (*38-40*). The SHP2 protein contains two SH2 domains (N-SH2/C-SH2), a catalytic PTP domain, and a C-terminal tail. SHP2 exists in the closed inactive conformation whereby the N-SH2 is wedged into the PTP domain, which is relieved upon pTyr binding on substrate proteins upon RTK signaling resulting in enzyme activity. Active SHP2 is marked by phosphorylation of Y546 and Y584 in the C-terminal tail (mostly referred to as Y542 and Y580 in literature based on the short isoform). We identified various phosphorylated tyrosine sites in SHP2, including Y546 and Y584. Phosphorylation at these sites was reduced in TKI-resistant cells, indicating SHP2 exists in the closed inactive state in the resistant cells. This may partly explain the paradoxically lower activation of RAS/MAPK signaling as evidenced by reduced phosphorylation of ERK and increased activation of the PI3K/AKT pathway as evidenced by increased phosphorylation of AKT in resistant cells (**Figure 5D**). Reduced SHP2 phosphorylation and activation may be partly a result of overall reduced activation of mutant EGFRs (**Figures 4 and 5**). However, further upstream inhibitors of SHP2 activity in TKI resistant cells remains to be elucidated.

The P100 assay includes a set of 100 phosphopeptide probes representing key cancer signaling pathways developed by the LINCS program (*41*). Phosphorylation of these sites from cell lines treated with a large set of therapeutic molecules was quantitated using a targeted MS-based P100 assay screen (*32*). Our anticorrelation analysis of quantitated phosphopeptides in our dataset that overlapped with the P100 assay peptides revealed dactolisib targeting PI3K/AKT and mTOR pathways as having potential to reverse TKI resistance. Dactolisib could circumvent osimertinib resistance both *in vitro* and *in vivo*. We were able to identify several other inhibitors of multiple signaling pathways that can potentially be used to treat TKI-resistant lung cancer that needs further validation. Our study underscored the possibility that the assay involving global or targeted MS-based phosphoproteomics can be used to identifying drug targets in the treatment of cancer.

In conclusion, this study identified and quantified, in an unbiased manner, the global proteome and phosphoproteome changes in 3^rd^ generation TKI resistant lung cancer cells in presence and absence of the respective TKIs. This study provides a new dimension and complements the existing studies involving genomics to identify resistance mechanisms and drug targets in only a few instances so far. Our study identified TKI resistant mechanisms and also an actionable target based on cellular signaling perturbations in resistant cells.

## Materials and Methods

### Cell culture and treatment

H1975 parental cell line was obtained from ATCC. The resistant cell lines AZR3/AZR4 (resistant to osimertinib) and COR1/COR10 (resistant to rociletinib) haave been described before (references) and were obtained from AstraZeneca and Colvis Oncology, respectively. The cells were cultured at 37 °C, 5% CO_2_ in RPMI medium 1640 (Pierce, Rockford, IL) with 10% dialyzed fetal bovine serum (Invitrogen, Carlsbad, CA) and 1% penicillin/streptomycin for at least five passages to completely label the proteome with stable isotope labeled heavy Arg and Lys for three-state SILAC experiments. The three state SILAC-labeled cells were TKI-sensitive cell line H1975 (light), the TKI-resistant cell lines AZR3 and COR1 (medium), and the TKI-resistant cell lines AZR4 and COR10 (heavy). The medium state cells were labeled with ^13^C_6_-Arg/D_4_-Lys and the heavy state cells were labeled with ^13^C_6_^15^N_4_-Arg/^13^C_6_^15^N_2_-Lys (Cambridge Iso-tope laboratories, Tewksbury, MA). Upon complete labeling, the cells were expanded to 15 cm dishes. The resistant cell lines were grown in the presence of TKIs until 3 days before the experiment. Each set of the SILAC labeled cells were treated either with DMSO or the corresponded drugs (50 nM osimertinib or 100 nM rociletinib) for one hour, respectively before harvesting.

### Protein extraction and sample processing

Cells were lysed with urea lysis buffer (20 mM HEPES pH 8.0, 8 M urea, 1 mM sodium orthovanadate, 2.5 mM sodium pyrophosphate and 1 mM-glycerophosphate). Protein concentrations were determined by the Modified Lowry method (BioRad, Hercules, CA). Equal amounts of protein from lysates of each SILAC state were mixed together. The combined lysate was reduced with 45 mM dithriothreitol (Sigma Aldrich, St. Louis, MO), alkylated with 100 mM iodoacetamide (Sigma Aldrich), and subsequently digested with trypsin (Worthington, NJ) at 37 °C overnight. The digest was then acidified with 1% TFA and peptides were desalted using C18 solid phase extraction columns (Supelco, Bellefonte, PA), lyophilized and stored at −80° C.

### Basic reversed phase liquid chromatography (RPLC) fractionation

Basic-RPLC separation was performed with an XBridge C18, 100 x 2.1 mm analytical column containing 5 μm particles and equipped with a 10 x 2.1 mm guard column (Waters, Milford, MA) with a flow rate of 0.25 ml/min. The solvent consisted of 10 mM triethylammonium bicarbonate (TEABC) as mobile phase A, and 10 mM TEABC in ACN as mobile phase B. Sample separation was accomplished using the following linear gradient: from 0 to 1% B in 5 min, from 1 to 10% B in 5 min, from 10 to 35% B in 30 min, and from 35 to 100% B in 5 min, and held at 100% B for an additional 3 min. A total of 96 fractions were collected during the LC separation in a 96-well plate containing 12.5 μl of 1% formic acid for immediate acidification. The collected fractions were concatenated into 12 fractions and dried in a vacuum centrifuge. 1/10^th^ of the peptides was injected directly for LC-MS/MS analysis from protein identification.

### Enrichment of Phosphopeptides

Remaining 9/10^th^ of the dried peptides were dissolved in solution A containing 80% ACN, 1% TFA, and 3% 2, 5-DHB and incubated with TiO_2_ (Titansphere, GL Sciences) pretreated (2 h at room temperature) with solution A. After 12 h, TiO_2_ beads were washed thrice using solution A and twice with 80% ACN containing 1% TFA. TiO_2_ bound peptides were eluted using 3% NH_4_OH in 40% ACN and immediately acidified using formic acid. The peptides were vacuum dried, C18 stage-tip cleaned before LC-MS/MS analysis.

Phosphotyrosine peptides were enriched using a PhosphoScan Kit (p-Tyr-100 and p-Tyr-1000, Cell Signaling, Danvers, MA). The lyophilized peptide was dissolved in IAP buffer (50 mM MOPS, pH 7.2, 10 mM sodium phosphate, 50 mM NaCl) and incubated with 40 μl of immobilized anti-phosphotyrosine antibody for 1 h at 4°C. The antibody beads were centrifuged for 1 min at 1500 x g, and the supernatant was separated and saved. The antibody-bound beads were washed 3 times with 1ml of IAP buffer and twice with water by inverting tube 5 times at 4 °C. The phosphotyrosine-containing peptides were eluted from antibody with 55 μl of 0.15% TFA by gently tapping the bottom of the tube and incubating at room temperature for 10 min. The peptides were vacuum dried, C18 stage-tip cleaned before LC-MS/MS analysis.

### LC-MS/MS Analysis

Peptides were analyzed on a Q Exactive HF interfaced with an Ultimate™ 3000 RSLCnano System (Thermo Fisher Scientific, San Jose, CA). The dried peptides and the enriched phosphopeptides were loaded onto a nano-trap column (Acclaim PepMap100 Nano Trap Column, C18, 5 μm, 100 Å, 100 μm i.d. x 2 cm) and separated on an Easy-spray™ C18 LC column (Acclaim PepMap100, C18, 2 μm, 100 Å, 75 μm i.d. x 25 cm). Mobile phases A and B consisted of 0.1% formic acid in water and 0.1% formic acid in 90% ACN, respectively. Peptides were eluted from the column at 300 nL/min using the following linear gradient: from 4 to 35% B in 60 min, from 35 to 45% B in 5 min, from 45 to 90% B in 5 min, and held at 90% B for an additional 5 min. The heated capillary temperature and spray voltage were 275 °C and 1.7 kV, respectively. Full spectra were collected from 350 to 1800 m/z in the Orbitrap analyzer at a resolution of 120,000, followed by data dependent HCD MS/MS scans of the fifteen most abundant ions at a resolution of 30,000, using 40% collision energy and dynamic exclusion time of 30s.

### Data Analysis

Peptides and proteins were identified and quantified using the MaxQuant software package (version 1.5.7.4) with the Andromeda search engine (*42, 43*). MS/MS spectra were searched against the Uniprot human protein database (Feb 2017, 70952 entries) and quantification was performed using default parameters for 3 state SILAC in MaxQuant. The parameters used for database search include trypsin as a protease with two missed cleavage sites allowed.

Carbamidomethyl cysteine was specified as a fixed modification. Phosphorylation at serine, threonine and tyrosine, deamidation of asparagine and glutamine, oxidation of methionine and protein N-terminal acetylation were specified as variable modifications. The precursor mass tolerance was set to 7 ppm and fragment mass tolerance to 20 ppm. False discovery rate was calculated using a decoy database and a 1% cut-off was applied to both peptide table and phosphosite table.

Combined normalized SILAC ratio of the proteins and the phosphopeptides and individual ratios of each experiment were obtained from the MaxQuant search. Perseus (version 1.5.5.3) was used to view and further analyze the data (*44*). Hierarchical clustering of proteins and phosphorylation were obtained in Perseus using log_2_ SILAC ratios. One sample t-test was performed in Perseus and the proteins or phosphopeptides with the p-value less than 0.05 and the SILAC ratio above 1.5-fold changes were considered as significantly changed.

### Kinase Enrichment Analysis

The goal of the enrichment analysis was to identify kinases whose targets were significantly over-represented among the phosphorylation sites that were significantly hyper- or hypo-phosphorylated in drug-resistant vs. sensitive cells when matched with the kinase-target site data in iPTMnet v.4.1 (*31*). To improve statistical power, the enrichment analysis was performed at the level of kinase families rather than individual kinases. Human kinases in iPTMnet were classified into families using a mapping table provided by KinBase (*45*)(http://kinase.com/web/current/human/; Dec 07 update). Phosphorylation sites that were significantly hyper- or hypo-phosphorylated in drug resistant vs. sensitive cells were selected using volcano plots (p-value greater than 0.05 and more than 1.5-fold change). Kinases for these sites were retrieved from iPTMnet using the iptmnetr R package (https://cran.r-project.org/web/packages/iptmnetr/index.html) and mapped to kinase families as above. An enrichment p-value for each family was obtained by performing Fisher’s Exact Test on the two-by-two contingency table consisting of: (i) the number of significant sites phosphorylated by at least one kinase in the family; (ii) the number of significant sites with at least one identified iPTMnet kinase that were not phosphorylated by any family members; (iii) the number of sites in iPTMnet phosphorylated by at least one kinase in the family; and (iv) the number of sites phosphorylated by at least one kinase in iPTMnet that were not phosphorylated by any family members. Only families whose kinases phosphorylated three or more significantly changed sites were included in the analysis. Calculations for significantly hyper- or hypophosphorylated sites were done separately. Multiple testing correction was performed with the Benjamin-Hochberg method. All statistical calculations were carried out using R.

### Drug Signature Comparison

Drug signature comparisons were performed using two methods. First, we used querying Touchstone-P, a library of phosphoproteomic signatures from several cell lines treated with a panel of small molecule drugs (*46*). Each signature in the library consists of the relative abundances of approximately 100 phosphorylation sites (P100). The Proteomic Query tool, available through the ConnectivityMap web interface (http://clue.io/proteomics-query), compares an input P100 phosphorylation signature to each signature in the library and reports a connectivity score ranging from −1 (strong negative connection/most “opposite” profile) to 1 (strong positive connection/most similar profile). The tool can calculate scores for data with and without replicates. To analyze our data with replicates, for each replicate of each cell line/drug combination in our dataset, we constructed a list of the log2 abundance ratios (TKI-resistant vs. sensitive) for all sites in the P100 set for which we had data. Replicates in which fewer than 40 of the P100 sites were observed were discarded. Depending on the cell line/drug combination, between two and four replicates had enough P100 data to be included in the analysis. The data was formatted as a gct file using the parse.gctx function in the cmap/cmapR R package (https://rdrr.io/github/cmap/cmapR) and submitted to the Proteomics Query tool to obtain connectivity scores. We also calculated the mean log2 abundance ratio across all replicates for the P100 sites observed in each cell line/drug combination in our data and submitted these profiles to the Proteomics Query tool using the “no replicates” option. Second, we calculated the Pearson correlation coefficient between each cell line/drug combination in our dataset and each experiment in the P100 dataset. We obtained the P100 data for this analysis from the P100 Level 3 data file P100Level3.08_16_2016.gct.txt (LDS-41234.tar.gz). Experiments in this dataset were performed in batches on 96-well plates. Abundance values for each P100 peptide were z-score normalized on a plate-by-plate basis. Values for multiple peptides representing the same site were merged by finding the median value; similarly, values from replicate experiments were merged by median. For each of our cell line/drug combinatins, we obtained the MaxQuant normalized ratios for each observed site, and selected that subset of sites that are present in the P100. Pearson correlations were calculated using R. Strongly negative correlations (closer to −1) indicate drugs that might reverse the resistance signature of the TKI-resistance cells.

### Immunoblot Analysis

For immunoblot analysis, cells were lysed in modified RIPA lysis buffer (150 mM NaCl, 1.0% IGEPAL CA-630, 0.5% sodium deoxy-cholate, and 50 mM Tris, pH 8.0), whereas for immunoprecipitation, cell extracts were prepared in Nonidet P-40 lysis buffer supplemented with protease and phosphatase inhibitors. 35 µg of lysate was separated by SDS-PAGE (4-15%, Invitrogen) and transferred to nitrocellulose membrane. After blocking in 5% BSA in PBS with 0.1% Tween 20 for 1 h, membranes were incubated with the appropriate primary antibody followed by secondary antibody coupled with HRP. The primary antibodies against phosphorylated and total proteins were purchased from Cell Signaling Technology (Andover, MA). Custom mouse monoclonal antibodies were made against EGFR L858R that recognizes mutant EGFR, but not wild-type EGFR, and against EGFR L858 that recognizes wild-type EGFR, but not mutant EGFR in collaboration with NanoTools (Germany).

### Drug Treatment and Cell Viability Assay

For drug treatment and cell survival analysis, cells were trypsinized with 0.25% trypsin in EDTA, and centrifuged at 1,000 rpm for 5 mins at room temperature. Cells were re-suspended in RPMI medium with 10% FBS, 1% pen/strep, counted with trypan blue exclusion and plated 4,000 cells per well in a 96 well-plate. Cells were allowed to settle overnight before drug treatment. 10x stock of dactolisib and VX970 was prepared fresh for each experiment. 10uL of drug was added to 90uL of RPMI media already present in each well. Cells were incubated at 37°C, 5% CO_2_ for 72 hours. After 72 hours of treatment, all medium was removed, and 50uL of 1x Promega CellTiter-Glo® Luminescent Cell Viability Assay reagent was added to each well and incubated at room temperature for 15 minutes. Luminescence was measured with a SpectraMax M5 microplate reader and recorded by SoftMax Pro 5.4.1. Raw luminescence was normalized and plotted with MS Excel.

### Quantitative RT-PCR

The All Prep DNA/RNA/Protein kit (QIAGEN) was used to total RNA from cell lines. The iScript Advanced cDNA Synthesis Kit (Bio-Rad) was used to synthesize cDNA to screen an Epithelial to Mesenchymal Transition (EMT) (Bio-Rad: SAB Target List) H384 plate using SsoAdvanced Universal SYBR Green Supermix (Bio-Rad). The plate was run on an ABI ViiA 7 instrument and analyzed with CFX manager software (Bio-Rad).

### Experimental Design and Statistical Rationale

Three and five biological replicates were processed for H1975/AZR3/AZR4 experiment and H1975/COR1/COR10 experiment, respectively. Both proteome and phosphoproteome data were acquired and the quantitation was done by 3-state SILAC labeling. Volcano plots were generated for the Log 2 ratio of M/L and H/L. Proteins or phosphosites with 1.5-fold change and p value greater than 0.05 were considered as significant.

## Acknowledgments

We thank AstraZeneca and Clovis Oncology for providing TKI resistant cell lines. We also thank Daren E. Cross (AstraZeneca) and Thomas Harding (Clovis Oncology) for helpful discussions. This research was supported by the NIH Intramural Research Program, Center of Cancer Research, National Cancer Institute, NIH U54HL127624 and NIH U01GM120953. Its contents are solely the responsibility of the authors and do not necessarily represent the official views of the NIH.

## Author Contributions

X.Z., T.M., and U.G. designed the research. X.Z., T.M., Y.Q., K.N., F.K., C.C. performed experiments. X.Z., T.M., K.R., J. C. and S.G. performed data analysis. C.W. and U.G. were involved in data analysis and discussion. X.Z. and U.G. wrote the manuscript. All authors reviewed, commented on, and approved the manuscript.

## Declaration of Interests

U.G. has a clinical trial agreement (CTA) with AstraZeneca for the current study and had received research funding from AstraZeneca, and Aurigene. U.G. is currently an employee of Bristol Myers Squibb. The other authors have no conflicts of interest to report.

## Data Availability

The MS proteomics data in this paper have been deposited in the ProteomeXchange Consortium (http://proteomecentral-.proteomexchange.org) via the PRIDE partner repository (*47, 48*) with the dataset identifier PXD020108. Username: Password: npddP7of

---

**Figure S1 – related to Figure 1.**
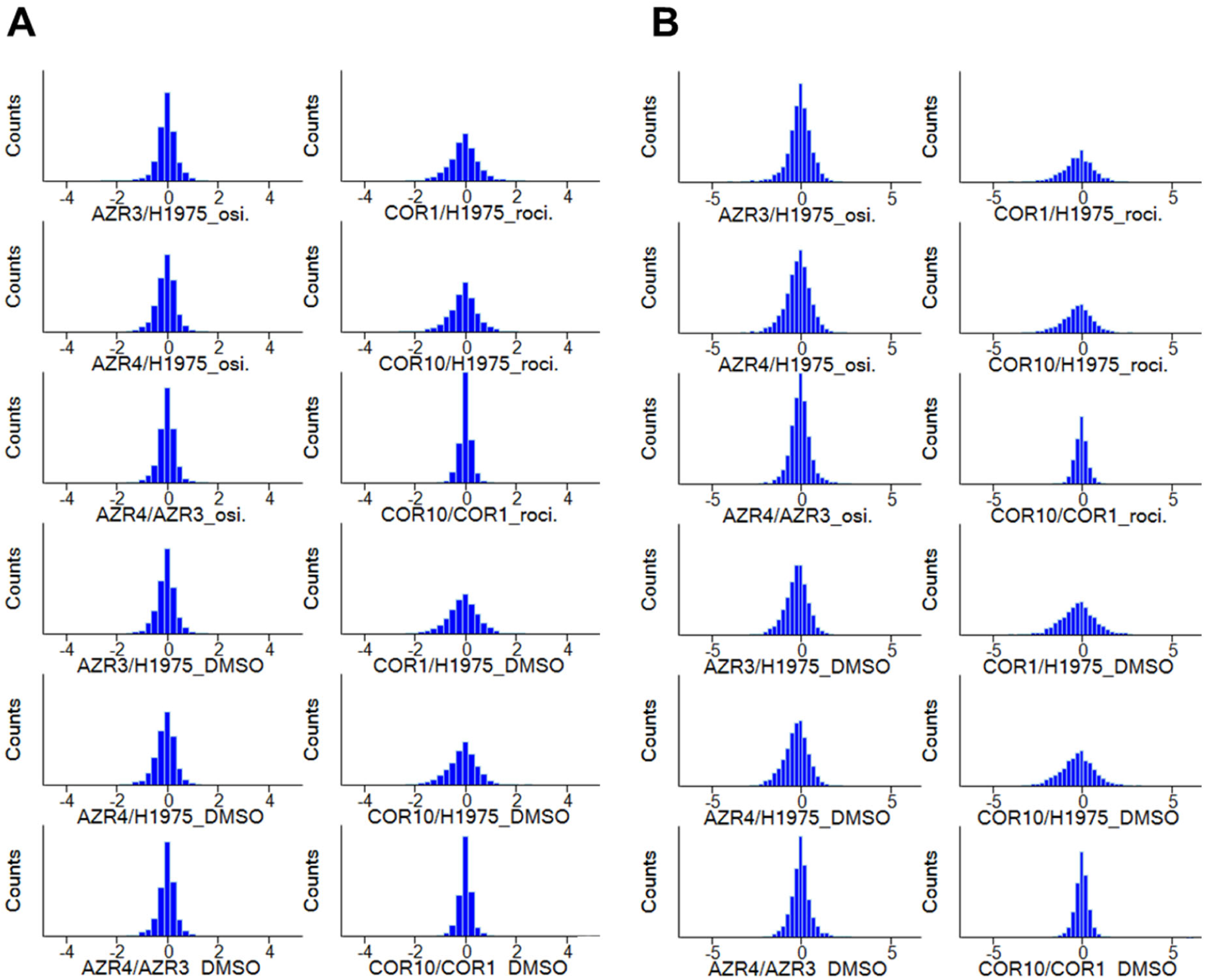
Histograms of the Log2 SILAC ratios between resistance and sensitive cell lines with DMSO or drug treatment of (A) total proteins and (B) phosphorylation sites.

**Figure S2 – related to Figure 1.**
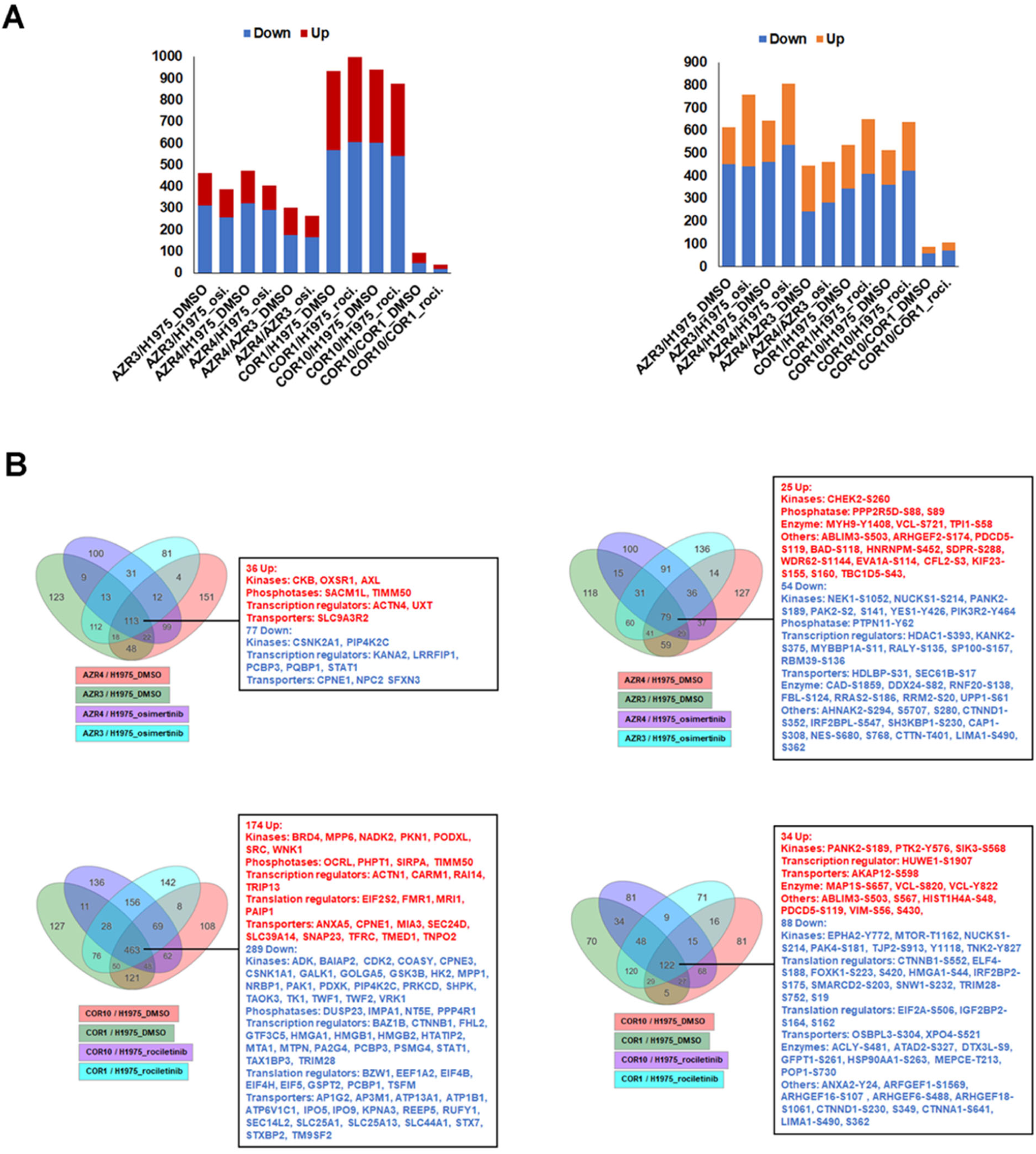
Significant alterations in protein and phosphopeptide abundance in TKI-sensitive and resistant cells. (A) Bar graphs represent number of proteins (left panel) and phosphopeptides (right panel) with statistically significant alteration in abundance between TKI-sensitive and resistant cells. (B) Venn diagrams showing the number of proteins (left) and phosphopeptides (right) whose abundance changed significantly between TKI-sensitive and resistant cells in presence or absence (DMSO) of corresponding TKI. Selected proteins or phosphopeptides with increased (red) or decreased (blue) abundance in the resistant cells are listed, that include kinases and phosphatases.

**Figure S3 – related to Figure 1 and Suppl. Fig. 2.**
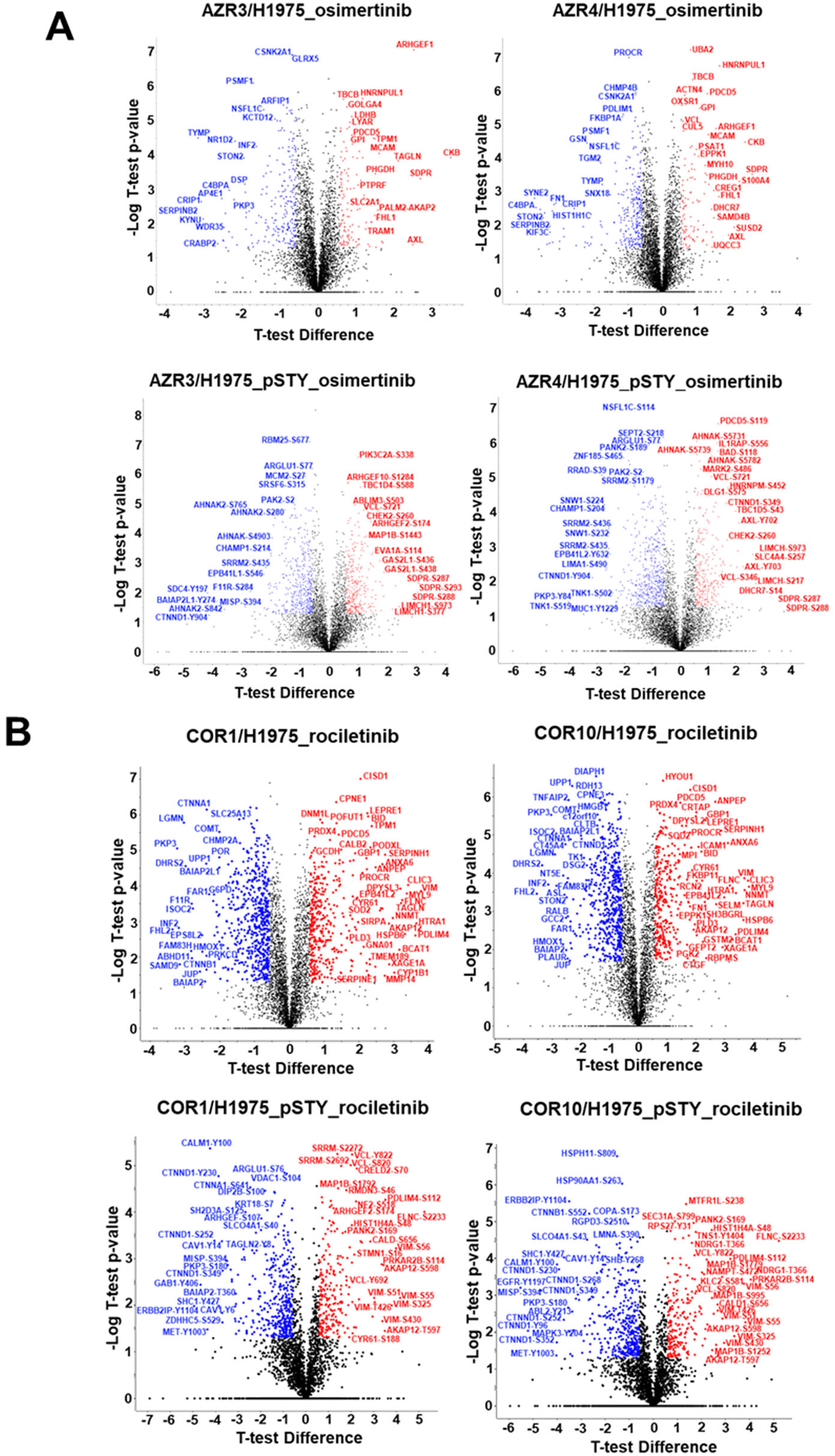
Volcano plots of the (A) total proteins and (B) phosphorylation sites between resistance and sensitive cell lines with drug treatment. X-axis is Log_2_ SILAC ratio; y-axis is – Log_10_ p-value. Proteins and phosphosites with significant regulation are highlighted in red (up) or blue (down).

**Figure S4 – related to Figure 2.**
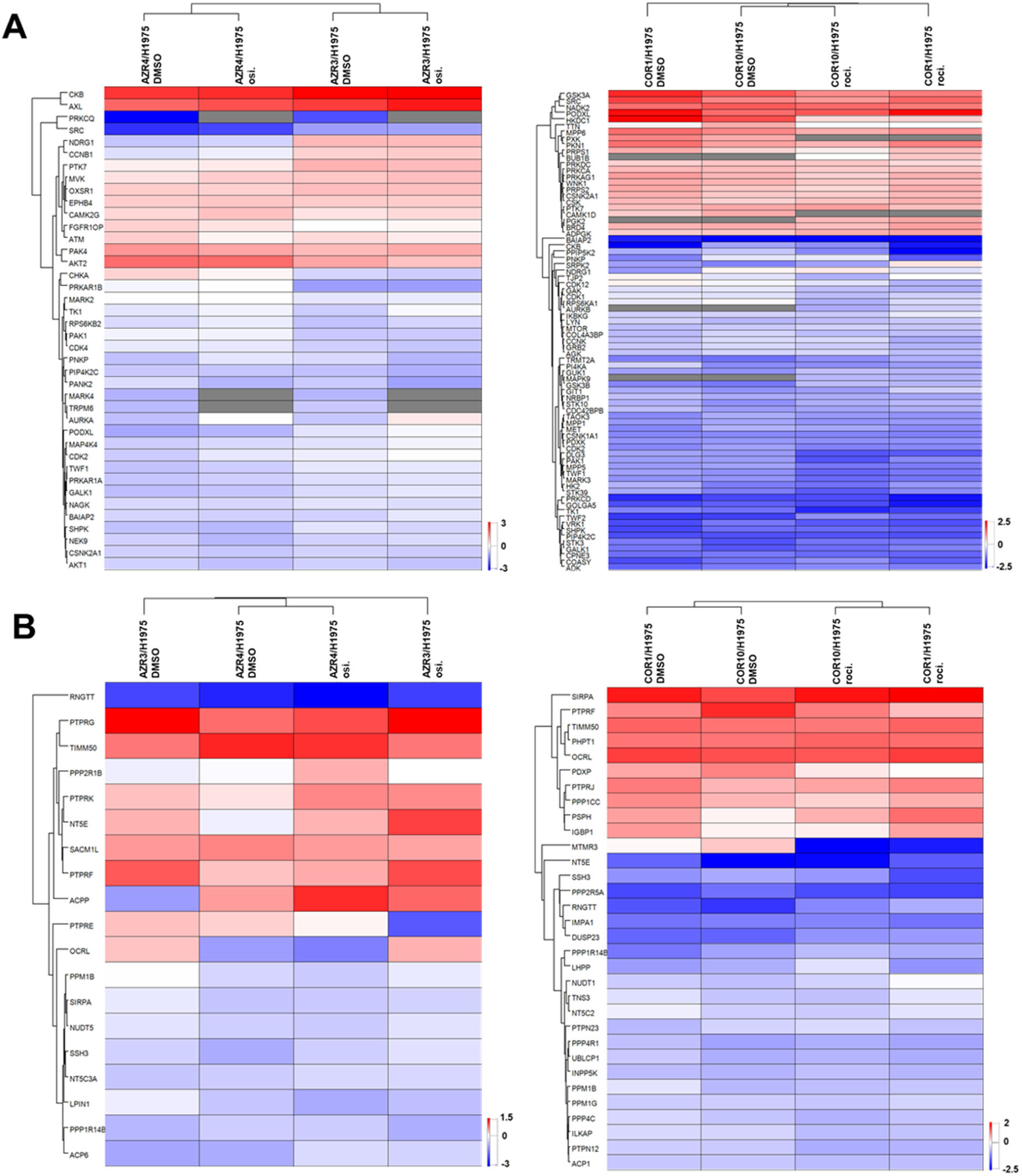
Hierarchical clustering of significantly altered abundances as reflected by the SILAC ratios (TKI-resistant/sensitive) of kinases (A) and phosphatases (B) in presence and absence (DMSO treatment) of respective TKI. (Left, osimertinib; Right, rociletinib). Columns represent different cell lines treated as indicated. Rows represent quantified kinases or phosphatases significantly altered in at least one column.

**Figure S5 – related to Figure 2.**
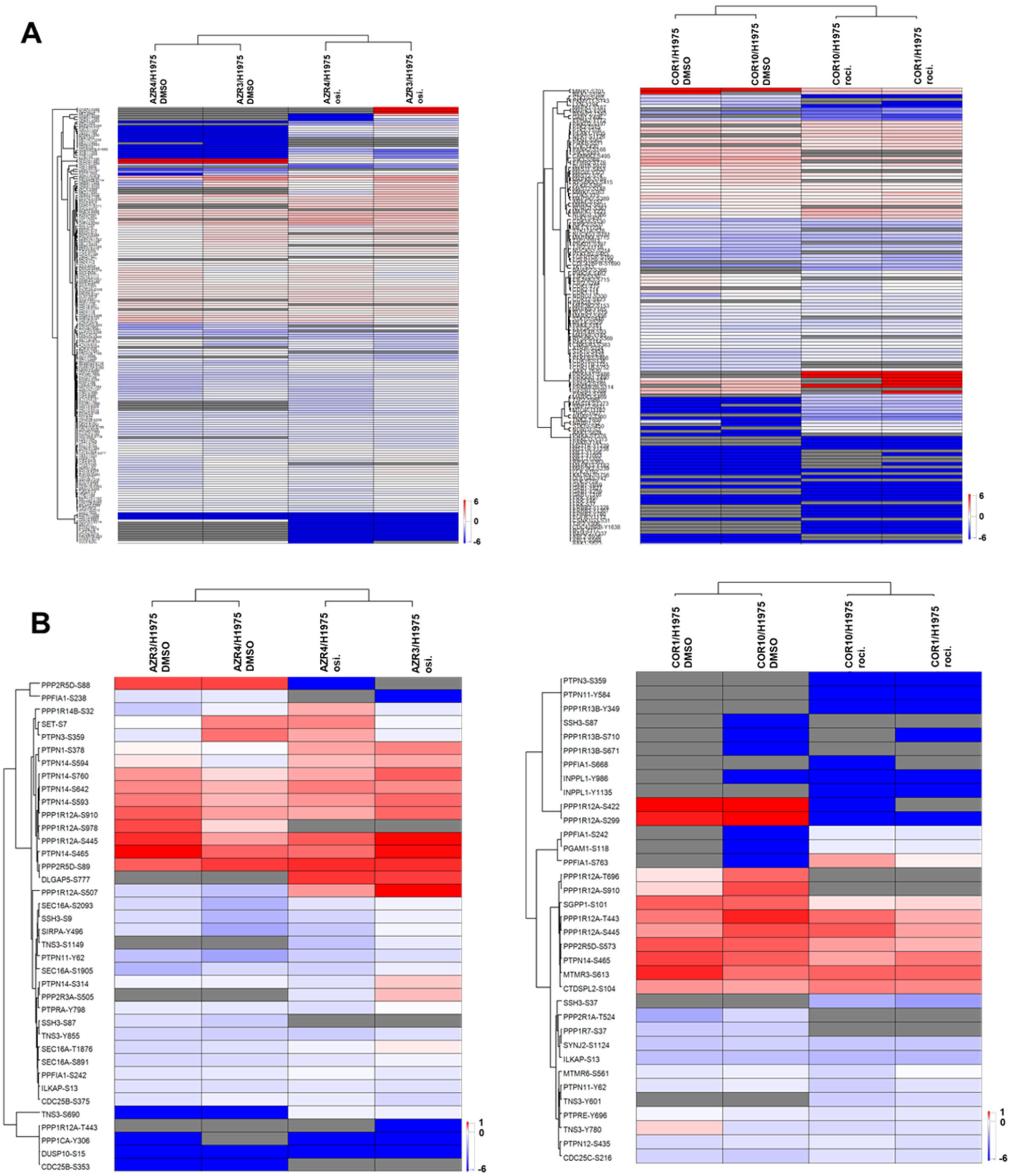
Hierarchical clustering of significantly altered phosphopeptides based on the SILAC ratios of phosphorylation (TKI-resistant/sensitive) of kinases (A) and phosphatases (B) in TKI-resistant cells with/without drug treatment. (Left, osimertinib; Right, rociletinib). Columns represent different cell lines treated as indicated. Rows represent quantified phosphosite containing phosphopeptides with significantly altered phosphorylation in at least one column.

**Figure S6 – related to Figure 4.**
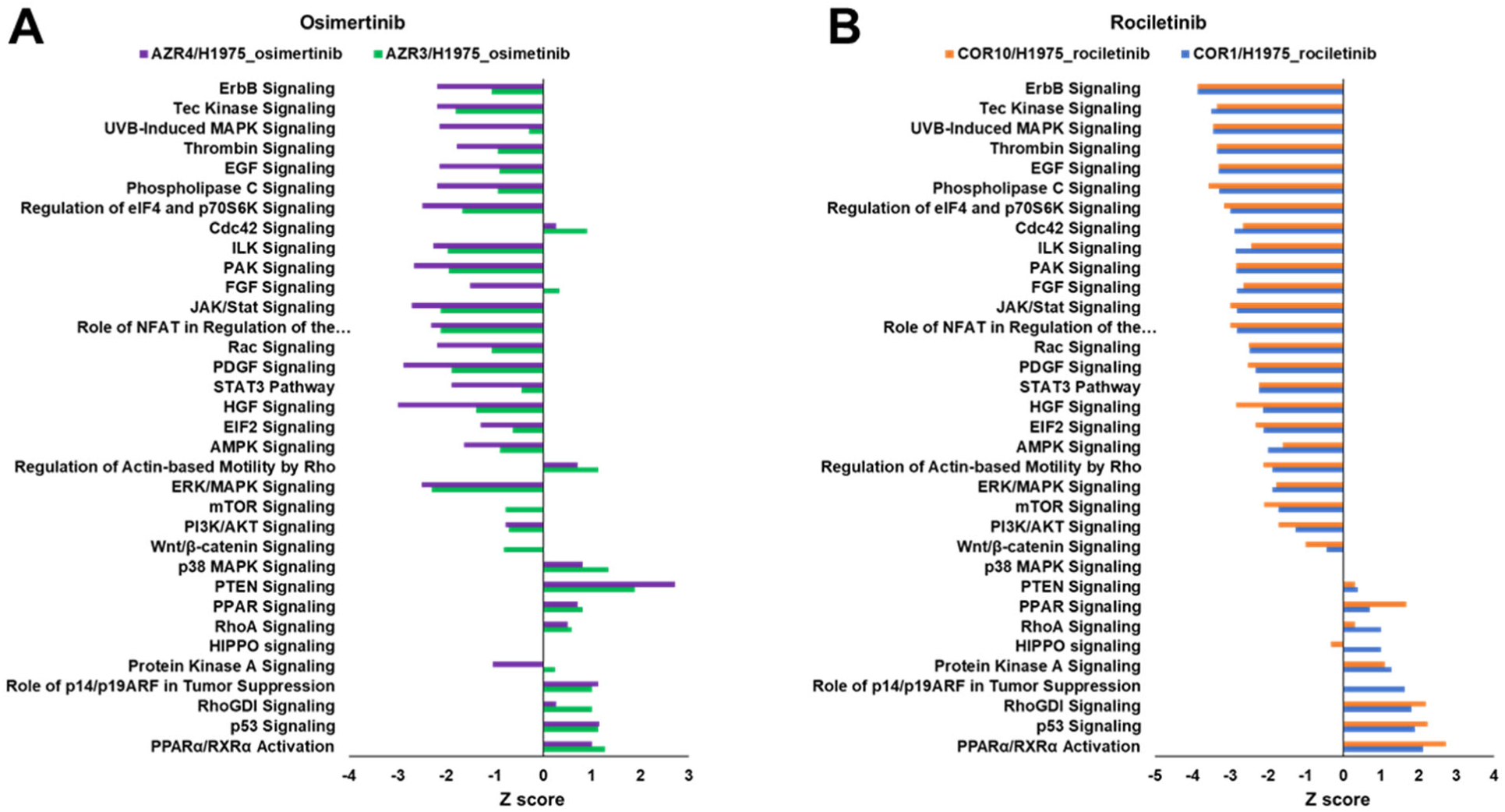
Canonical pathways enriched in (A) osimertinib and (B) rociletinib resistant cells, identified using IPA analysis of genes with significantly altered phosphorylation in specific phosphopeptides.

**Figure S7 – related to Figure 5.**
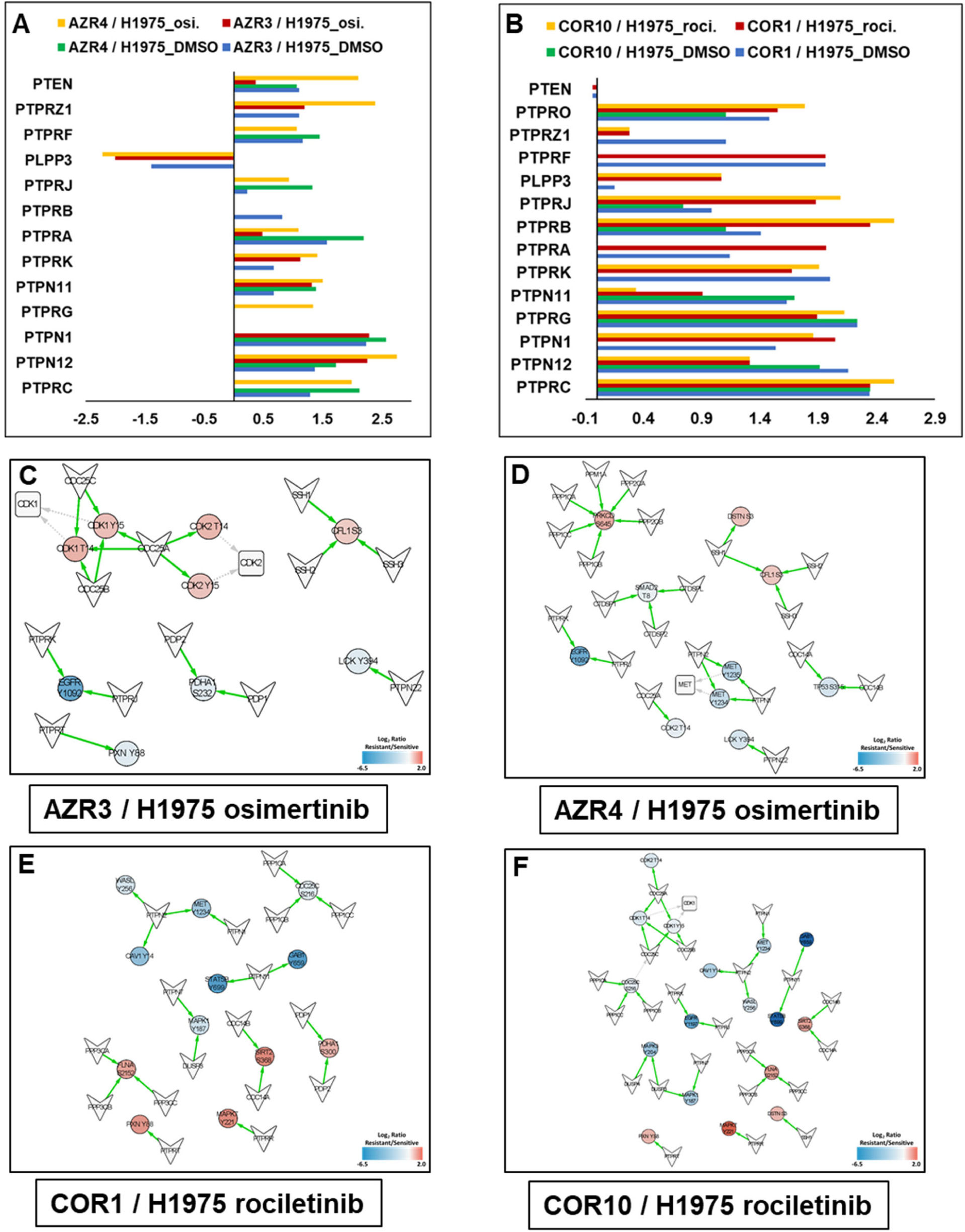
Potential phosphatases upstream of differentially phosphorylated targets in TKI-resistant cells. Upstream phosphatases of significantly altered phosphopeptide substrates identified using IPA analysis in (A) osimertinib and (B) rociletinib resistant cells. (C) Network connecting phosphosites that are significantly altered in TKI-resistant cells to their experimentally validated phosphatases obtained from the human DEPhOsphorylation Database (DEPOD).

## Notes

### Competing Interest Statement

The authors have declared no competing interest.

## References

1. R. Siegel, D. Naishadham, A. Jemal, Cancer statistics, 2013. CA Cancer J Clin 63, 11–30 (2013); published online EpubJan (10.3322/caac.21166).

2. W. Pao, V. A. Miller, K. A. Politi, G. J. Riely, R. Somwar, M. F. Zakowski, M. G. Kris, H. Varmus, Acquired resistance of lung adenocarcinomas to gefitinib or erlotinib is associated with a second mutation in the EGFR kinase domain. PLoS Med 2, e73 (2005); published online EpubMar (10.1371/journal.pmed.0020073).

3. D. A. Cross, S. E. Ashton, S. Ghiorghiu, C. Eberlein, C. A. Nebhan, P. J. Spitzler, J. P. Orme, M. R. Finlay, R. A. Ward, M. J. Mellor, G. Hughes, A. Rahi, V. N. Jacobs, M. Red Brewer, E. Ichihara, J. Sun, H. Jin, P. Ballard, K. Al-Kadhimi, R. Rowlinson, T. Klinowska, G. H. Richmond, M. Cantarini, D. W. Kim, M. R. Ranson, W. Pao, AZD9291, an irreversible EGFR TKI, overcomes T790M-mediated resistance to EGFR inhibitors in lung cancer. Cancer Discov 4, 1046–1061 (2014); published online EpubSep (10.1158/2159-8290.CD-14-0337).

4. J. C. Soria, Y. Ohe, J. Vansteenkiste, T. Reungwetwattana, B. Chewaskulyong, K. H. Lee, A. Dechaphunkul, F. Imamura, N. Nogami, T. Kurata, I. Okamoto, C. Zhou, B. C. Cho, Y. Cheng, E. K. Cho, P. J. Voon, D. Planchard, W. C. Su, J. E. Gray, S. M. Lee, R. Hodge, M. Marotti, Y. Rukazenkov, S. S. Ramalingam, F. Investigators, Osimertinib in Untreated EGFR-Mutated Advanced Non-Small-Cell Lung Cancer. N Engl J Med 378, 113–125 (2018); published online EpubJan 11 (10.1056/NEJMoa1713137).

5. P. A. Janne, J. C. Yang, D. W. Kim, D. Planchard, Y. Ohe, S. S. Ramalingam, M. J. Ahn, S. W. Kim, W. C. Su, L. Horn, D. Haggstrom, E. Felip, J. H. Kim, P. Frewer, M. Cantarini, K. H. Brown, P. A. Dickinson, S. Ghiorghiu, M. Ranson, AZD9291 in EGFR inhibitor-resistant non-small-cell lung cancer. N Engl J Med 372, 1689–1699 (2015); published online EpubApr 30 (10.1056/NEJMoa1411817).

6. L. V. Sequist, J. C. Soria, J. W. Goldman, H. A. Wakelee, S. M. Gadgeel, A. Varga, V. Papadimitrakopoulou, B. J. Solomon, G. R. Oxnard, R. Dziadziuszko, D. L. Aisner, R. C. Doebele, C. Galasso, E. B. Garon, R. S. Heist, J. Logan, J. W. Neal, M. A. Mendenhall, S. Nichols, Z. Piotrowska, A. J. Wozniak, M. Raponi, C. A. Karlovich, S. Jaw-Tsai, J. Isaacson, D. Despain, S. L. Matheny, L. Rolfe, A. R. Allen, D. R. Camidge, Rociletinib in EGFR-mutated non-small-cell lung cancer. N Engl J Med 372, 1700–1709 (2015); published online EpubApr 30 (10.1056/NEJMoa1413654).

7. L. V. Sequist, J. C. Soria, D. R. Camidge, Update to Rociletinib Data with the RECIST Confirmed Response Rate. N Engl J Med 374, 2296–2297 (2016); published online EpubJun 9 (10.1056/NEJMc1602688).

8. L. V. Sequist, Z. Piotrowska, M. J. Niederst, R. S. Heist, S. Digumarthy, A. T. Shaw, J. A. Engelman, Osimertinib Responses After Disease Progression in Patients Who Had Been Receiving Rociletinib. JAMA Oncol 2, 541–543 (2016); published online EpubApr (10.1001/jamaoncol.2015.5009).

9. K. S. Thress, C. P. Paweletz, E. Felip, B. C. Cho, D. Stetson, B. Dougherty, Z. Lai, A. Markovets, A. Vivancos, Y. Kuang, D. Ercan, S. E. Matthews, M. Cantarini, J. C. Barrett, P. A. Janne, G. R. Oxnard, Acquired EGFR C797S mutation mediates resistance to AZD9291 in non-small cell lung cancer harboring EGFR T790M. Nat Med 21, 560–562 (2015); published online EpubJun (10.1038/nm.3854).

10. Z. Piotrowska, M. J. Niederst, C. A. Karlovich, H. A. Wakelee, J. W. Neal, M. Mino-Kenudson, L. Fulton, A. N. Hata, E. L. Lockerman, A. Kalsy, S. Digumarthy, A. Muzikansky, M. Raponi, A. R. Garcia, H. E. Mulvey, M. K. Parks, R. H. DiCecca, D. Dias-Santagata, A. J. Iafrate, A. T. Shaw, A. R. Allen, J. A. Engelman, L. V. Sequist, Heterogeneity Underlies the Emergence of EGFRT790 Wild-Type Clones Following Treatment of T790M-Positive Cancers with a Third-Generation EGFR Inhibitor. Cancer Discov 5, 713–722 (2015); published online EpubJul (10.1158/2159-8290.CD-15-0399).

11. N. Roper, A. L. Brown, J. S. Wei, S. Pack, C. Trindade, C. Kim, O. Restifo, S. Gao, S. Sindiri, F. Mehrabadi, R. El Meskini, Z. W. Ohler, T. K. Maity, A. Venugopalan, C. M. Cultraro, E. Akoth, E. Padiernos, H. Chen, A. Kesarwala, D. K. Smart, N. Nilubol, A. Rajan, Z. Piotrowska, L. Xi, M. Raffeld, A. R. Panchenko, C. Sahinalp, S. Hewitt, C. D. Hoang, J. Khan, U. Guha, Clonal Evolution and Heterogeneity of Osimertinib Acquired Resistance Mechanisms in EGFR Mutant Lung Cancer. Cell Rep Med 1, (2020); published online EpubApr 21 (10.1016/j.xcrm.2020.100007).

12. G. R. Oxnard, Y. Hu, K. F. Mileham, H. Husain, D. B. Costa, P. Tracy, N. Feeney, L. M. Sholl, S. E. Dahlberg, A. J. Redig, D. J. Kwiatkowski, M. S. Rabin, C. P. Paweletz, K. S. Thress, P. A. Janne, Assessment of Resistance Mechanisms and Clinical Implications in Patients With EGFR T790M-Positive Lung Cancer and Acquired Resistance to Osimertinib. JAMA Oncol 4, 1527–1534 (2018); published online EpubNov 1 (10.1001/jamaoncol.2018.2969).

13. X. Le, S. Puri, M. V. Negrao, M. B. Nilsson, J. Robichaux, T. Boyle, J. K. Hicks, K. L. Lovinger, E. Roarty, W. Rinsurongkawong, M. Tang, H. Sun, Y. Elamin, L. C. Lacerda, J. Lewis, J. A. Roth, S. G. Swisher, J. J. Lee, W. N. William, Jr., B. S. Glisson, J. Zhang, V. A. Papadimitrakopoulou, J. E. Gray, J. V. Heymach, Landscape of EGFR-Dependent and -Independent Resistance Mechanisms to Osimertinib and Continuation Therapy Beyond Progression in EGFR-Mutant NSCLC. Clin Cancer Res 24, 6195–6203 (2018); published online EpubDec 15 (10.1158/1078-0432.CCR-18-1542).

14. Z. Yang, N. Yang, Q. Ou, Y. Xiang, T. Jiang, X. Wu, H. Bao, X. Tong, X. Wang, Y. W. Shao, Y. Liu, Y. Wang, C. Zhou, Investigating Novel Resistance Mechanisms to Third-Generation EGFR Tyrosine Kinase Inhibitor Osimertinib in Non-Small Cell Lung Cancer Patients. Clin Cancer Res 24, 3097–3107 (2018); published online EpubJul 1 (10.1158/1078-0432.CCR-17-2310).

15. Z. Piotrowska, H. Isozaki, J. K. Lennerz, J. F. Gainor, I. T. Lennes, V. W. Zhu, N. Marcoux, M. K. Banwait, S. R. Digumarthy, W. Su, S. Yoda, A. K. Riley, V. Nangia, J. J. Lin, R. J. Nagy, R. B. Lanman, D. Dias-Santagata, M. Mino-Kenudson, A. J. Iafrate, R. S. Heist, A. T. Shaw, E. K. Evans, C. Clifford, S. I. Ou, B. Wolf, A. N. Hata, L. V. Sequist, Landscape of Acquired Resistance to Osimertinib in EGFR-Mutant NSCLC and Clinical Validation of Combined EGFR and RET Inhibition with Osimertinib and BLU-667 for Acquired RET Fusion. Cancer Discov 8, 1529–1539 (2018); published online EpubDec (10.1158/2159-8290.CD-18-1022).

16. Z. H. Tang, J. J. Lu, Osimertinib resistance in non-small cell lung cancer: Mechanisms and therapeutic strategies. Cancer Lett 420, 242–246 (2018); published online EpubApr 28 (10.1016/j.canlet.2018.02.004).

17. A. Murtuza, A. Bulbul, J. P. Shen, P. Keshavarzian, B. D. Woodward, F. J. Lopez-Diaz, S. M. Lippman, H. Husain, Novel Third-Generation EGFR Tyrosine Kinase Inhibitors and Strategies to Overcome Therapeutic Resistance in Lung Cancer. Cancer Res 79, 689–698 (2019); published online EpubFeb 15 (10.1158/0008-5472.CAN-18-1281).

18. X. Zhang, N. Belkina, H. K. Jacob, T. Maity, R. Biswas, A. Venugopalan, P. G. Shaw, M. S. Kim, R. Chaerkady, A. Pandey, U. Guha, Identifying novel targets of oncogenic EGF receptor signaling in lung cancer through global phosphoproteomics. Proteomics 15, 340–355 (2015); published online EpubJan (10.1002/pmic.201400315).

19. X. Zhang, T. Maity, M. K. Kashyap, M. Bansal, A. Venugopalan, S. Singh, S. Awasthi, A. Marimuthu, H. K. Charles Jacob, N. Belkina, S. Pitts, C. M. Cultraro, S. Gao, G. Kirkali, R. Biswas, R. Chaerkady, A. Califano, A. Pandey, U. Guha, Quantitative Tyrosine Phosphoproteomics of Epidermal Growth Factor Receptor (EGFR) Tyrosine Kinase Inhibitor-treated Lung Adenocarcinoma Cells Reveals Potential Novel Biomarkers of Therapeutic Response. Mol Cell Proteomics 16, 891–910 (2017); published online EpubMay (10.1074/mcp.M117.067439).

20. Z. Zhang, J. C. Lee, L. Lin, V. Olivas, V. Au, T. LaFramboise, M. Abdel-Rahman, X. Wang, A. D. Levine, J. K. Rho, Y. J. Choi, C. M. Choi, S. W. Kim, S. J. Jang, Y. S. Park, W. S. Kim, D. H. Lee, J. S. Lee, V. A. Miller, M. Arcila, M. Ladanyi, P. Moonsamy, C. Sawyers, T. J. Boggon, P. C. Ma, C. Costa, M. Taron, R. Rosell, B. Halmos, T. G. Bivona, Activation of the AXL kinase causes resistance to EGFR-targeted therapy in lung cancer. Nat Genet 44, 852–860 (2012); published online EpubJul 1 (10.1038/ng.2330).

21. L. A. Byers, L. Diao, J. Wang, P. Saintigny, L. Girard, M. Peyton, L. Shen, Y. Fan, U. Giri, P. K. Tumula, M. B. Nilsson, J. Gudikote, H. Tran, R. J. Cardnell, D. J. Bearss, S. L. Warner, J. M. Foulks, S. B. Kanner, V. Gandhi, N. Krett, S. T. Rosen, E. S. Kim, R. S. Herbst, G. R. Blumenschein, J. J. Lee, S. M. Lippman, K. K. Ang, G. B. Mills, W. K. Hong, J. N. Weinstein, Wistuba, II, K. R. Coombes, J. D. Minna, J. V. Heymach, An epithelial-mesenchymal transition gene signature predicts resistance to EGFR and PI3K inhibitors and identifies Axl as a therapeutic target for overcoming EGFR inhibitor resistance. Clin Cancer Res 19, 279–290 (2013); published online EpubJan 1 (10.1158/1078-0432.CCR-12-1558).

22. E. Izumchenko, X. Chang, C. Michailidi, L. Kagohara, R. Ravi, K. Paz, M. Brait, M. O. Hoque, S. Ling, A. Bedi, D. Sidransky, The TGFbeta-miR200-MIG6 pathway orchestrates the EMT-associated kinase switch that induces resistance to EGFR inhibitors. Cancer Res 74, 3995–4005 (2014); published online EpubJul 15 (10.1158/0008-5472.CAN-14-0110).

23. R. L. Yauch, T. Januario, D. A. Eberhard, G. Cavet, W. Zhu, L. Fu, T. Q. Pham, R. Soriano, J. Stinson, S. Seshagiri, Z. Modrusan, C. Y. Lin, V. O’Neill, L. C. Amler, Epithelial versus mesenchymal phenotype determines in vitro sensitivity and predicts clinical activity of erlotinib in lung cancer patients. Clin Cancer Res 11, 8686–8698 (2005); published online EpubDec 15 (10.1158/1078-0432.CCR-05-1492).

24. N. Bin-Nun, H. Lichtig, A. Malyarova, M. Levy, S. Elias, D. Frank, PTK7 modulates Wnt signaling activity via LRP6. Development 141, 410–421 (2014); published online EpubJan (10.1242/dev.095984).

25. N. R. Dunn, N. S. Tolwinski, Ptk7 and Mcc, Unfancied Components in Non-Canonical Wnt Signaling and Cancer. Cancers (Basel) 8, (2016); published online EpubJul 16 (10.3390/cancers8070068).

26. M. Hayes, M. Naito, A. Daulat, S. Angers, B. Ciruna, Ptk7 promotes non-canonical Wnt/PCP-mediated morphogenesis and inhibits Wnt/beta-catenin-dependent cell fate decisions during vertebrate development. Development 140, 1807–1818 (2013); published online EpubApr (10.1242/dev.090183).

27. H. Peradziryi, N. A. Kaplan, M. Podleschny, X. Liu, P. Wehner, A. Borchers, N. S. Tolwinski, PTK7/Otk interacts with Wnts and inhibits canonical Wnt signalling. EMBO J 30, 3729–3740 (2011); published online EpubJul 19 (10.1038/emboj.2011.236).

28. S. Li, J. Cao, W. Zhang, F. Zhang, G. Ni, Q. Luo, M. Wang, X. Tao, H. Xia, Protein tyrosine phosphatase PTPN3 promotes drug resistance and stem cell-like characteristics in ovarian cancer. Sci Rep 6, 36873 (2016); published online EpubNov 11 (10.1038/srep36873).

29. A. S. Kumar, I. Naruszewicz, P. Wang, C. Leung-Hagesteijn, G. E. Hannigan, ILKAP regulates ILK signaling and inhibits anchorage-independent growth. Oncogene 23, 3454–3461 (2004); published online EpubApr 22 (10.1038/sj.onc.1207473).

30. N. P. Damle, M. Kohn, The human DEPhOsphorylation Database DEPOD: 2019 update. Database (Oxford) 2019, (2019); published online EpubJan 1 (10.1093/database/baz133).

31. H. Huang, C. N. Arighi, K. E. Ross, J. Ren, G. Li, S. C. Chen, Q. Wang, J. Cowart, K. Vijay-Shanker, C. H. Wu, iPTMnet: an integrated resource for protein post-translational modification network discovery. Nucleic Acids Res 46, D542–D550 (2018); published online EpubJan 4 (10.1093/nar/gkx1104).

32. J. G. Abelin, J. Patel, X. Lu, C. M. Feeney, L. Fagbami, A. L. Creech, R. Hu, D. Lam, D. Davison, L. Pino, J. W. Qiao, E. Kuhn, A. Officer, J. Li, S. Abbatiello, A. Subramanian, R. Sidman, E. Snyder, S. A. Carr, J. D. Jaffe, Reduced-representation Phosphosignatures Measured by Quantitative Targeted MS Capture Cellular States and Enable Large-scale Comparison of Drug-induced Phenotypes. Mol Cell Proteomics 15, 1622–1641 (2016); published online EpubMay (10.1074/mcp.M116.058354).

33. A. O. Walter, R. T. Sjin, H. J. Haringsma, K. Ohashi, J. Sun, K. Lee, A. Dubrovskiy, M. Labenski, Z. Zhu, Z. Wang, M. Sheets, T. St Martin, R. Karp, D. van Kalken, P. Chaturvedi, D. Niu, M. Nacht, R. C. Petter, W. Westlin, K. Lin, S. Jaw-Tsai, M. Raponi, T. Van Dyke, J. Etter, Z. Weaver, W. Pao, J. Singh, A. D. Simmons, T. C. Harding, A. Allen, Discovery of a mutant-selective covalent inhibitor of EGFR that overcomes T790M-mediated resistance in NSCLC. Cancer Discov 3, 1404–1415 (2013); published online EpubDec (10.1158/2159-8290.CD-13-0314).

34. M. Grzmil, B. A. Hemmings, Translation regulation as a therapeutic target in cancer. Cancer Res 72, 3891–3900 (2012); published online EpubAug 15 (10.1158/0008-5472.CAN-12-0026).

35. L. Zhao, J. R. Whiteaker, M. E. Pope, E. Kuhn, A. Jackson, N. L. Anderson, T. W. Pearson, S. A. Carr, A. G. Paulovich, Quantification of proteins using peptide immunoaffinity enrichment coupled with mass spectrometry. J Vis Exp, (2011); published online EpubJul 31 (10.3791/2812).

36. J. R. Whiteaker, L. Zhao, S. E. Abbatiello, M. Burgess, E. Kuhn, C. Lin, M. E. Pope, M. Razavi, N. L. Anderson, T. W. Pearson, S. A. Carr, A. G. Paulovich, Evaluation of large scale quantitative proteomic assay development using peptide affinity-based mass spectrometry. Mol Cell Proteomics 10, M110 005645 (2011); published online EpubApr (10.1074/mcp.M110.005645).

37. J. R. Whiteaker, L. Zhao, L. Anderson, A. G. Paulovich, An automated and multiplexed method for high throughput peptide immunoaffinity enrichment and multiple reaction monitoring mass spectrometry-based quantification of protein biomarkers. Mol Cell Proteomics 9, 184–196 (2010); published online EpubJan (10.1074/mcp.M900254-MCP200).

38. M. Tajan, A. de Rocca Serra, P. Valet, T. Edouard, A. Yart, SHP2 sails from physiology to pathology. Eur J Med Genet 58, 509–525 (2015); published online EpubOct (10.1016/j.ejmg.2015.08.005).

39. S. Q. Zhang, W. G. Tsiaras, T. Araki, G. Wen, L. Minichiello, R. Klein, B. G. Neel, Receptor-specific regulation of phosphatidylinositol 3’-kinase activation by the protein tyrosine phosphatase Shp2. Mol Cell Biol 22, 4062–4072 (2002); published online EpubJun (10.1128/mcb.22.12.4062-4072.2002).

40. A. Montagner, A. Yart, M. Dance, B. Perret, J. P. Salles, P. Raynal, A novel role for Gab1 and SHP2 in epidermal growth factor-induced Ras activation. J Biol Chem 280, 5350–5360 (2005); published online EpubFeb 18 (10.1074/jbc.M410012200).

41. V. Stathias, J. Turner, A. Koleti, D. Vidovic, D. Cooper, M. Fazel-Najafabadi, M. Pilarczyk, R. Terryn, C. Chung, A. Umeano, D. J. B. Clarke, A. Lachmann, J. E. Evangelista, A. Ma’ayan, M. Medvedovic, S. C. Schurer, LINCS Data Portal 2.0: next generation access point for perturbation-response signatures. Nucleic Acids Res 48, D431–D439 (2020); published online EpubJan 8 (10.1093/nar/gkz1023).

42. J. Cox, M. Mann, MaxQuant enables high peptide identification rates, individualized p.p.b.-range mass accuracies and proteome-wide protein quantification. Nat Biotechnol 26, 1367–1372 (2008); published online EpubDec (10.1038/nbt.1511).

43. J. Cox, N. Neuhauser, A. Michalski, R. A. Scheltema, J. V. Olsen, M. Mann, Andromeda: a peptide search engine integrated into the MaxQuant environment. J Proteome Res 10, 1794–1805 (2011); published online EpubApr 1 (10.1021/pr101065j).

44. S. Tyanova, J. Cox, Perseus: A Bioinformatics Platform for Integrative Analysis of Proteomics Data in Cancer Research. Methods Mol Biol 1711, 133–148 (2018)10.1007/978-1-4939-7493-1_7).

45. G. Manning, D. B. Whyte, R. Martinez, T. Hunter, S. Sudarsanam, The protein kinase complement of the human genome. Science 298, 1912–1934 (2002); published online EpubDec 6 (10.1126/science.1075762).

46. L. Litichevskiy, R. Peckner, J. G. Abelin, J. K. Asiedu, A. L. Creech, J. F. Davis, D. Davison, C. M. Dunning, J. D. Egertson, S. Egri, J. Gould, T. Ko, S. A. Johnson, D. L. Lahr, D. Lam, Z. Liu, N. J. Lyons, X. Lu, B. X. MacLean, A. E. Mungenast, A. Officer, T. E. Natoli, M. Papanastasiou, J. Patel, V. Sharma, C. Toder, A. A. Tubelli, J. Z. Young, S. A. Carr, T. R. Golub, A. Subramanian, M. J. MacCoss, L. H. Tsai, J. D. Jaffe, A Library of Phosphoproteomic and Chromatin Signatures for Characterizing Cellular Responses to Drug Perturbations. Cell Syst 6, 424–443 e427 (2018); published online EpubApr 25 (10.1016/j.cels.2018.03.012).

47. J. A. Vizcaino, R. G. Cote, A. Csordas, J. A. Dianes, A. Fabregat, J. M. Foster, J. Griss, E. Alpi, M. Birim, J. Contell, G. O’Kelly, A. Schoenegger, D. Ovelleiro, Y. Perez-Riverol, F. Reisinger, D. Rios, R. Wang, H. Hermjakob, The PRoteomics IDEntifications (PRIDE) database and associated tools: status in 2013. Nucleic Acids Res 41, D1063–1069 (2013); published online EpubJan (10.1093/nar/gks1262).

48. J. A. Vizcaino, A. Csordas, N. del-Toro, J. A. Dianes, J. Griss, I. Lavidas, G. Mayer, Y. Perez-Riverol, F. Reisinger, T. Ternent, Q. W. Xu, R. Wang, H. Hermjakob, 2016 update of the PRIDE database and its related tools. Nucleic Acids Res 44, D447–456 (2016); published online EpubJan 4 (10.1093/nar/gkv1145).

